# Automated Synthetic Cell-based Screening for Designed Proteins with Emergent Functions

**DOI:** 10.64898/2026.05.07.723257

**Authors:** Kareem Al Nahas, Béla P. Frohn, Aleksandra Šakanović, Frank Siedler, Petra Schwille

## Abstract

Designing minimal biological systems with emergent functions such as spatiotemporal self-organization is a central goal of bottom-up synthetic biology. While computational optimization and design show promises to accelerate functional protein engineering through Design-Build-Test-Learn cycles, screening libraries for complex functions remains a major challenge. Conventional screens typically lack the spatiotemporal resolution and cell-like confinement required in bottom-up synthetic biology. Here, we present PUREdrop, an automated microfluidic platform that encapsulates and expresses protein libraries in thousands of picolitre-sized synthetic cells per construct. These droplets are sorted into a 96-well plate and analyzed by time-lapse imaging, allowing parallel quantification of expression kinetics and emergent functions. To demonstrate the platform’s potential, we first screened computationally re-designed variants of the bacterial cell division protein FtsZ, identifying variants with improved bundling phenotypes and faster kinetics. We then extended our screening procedure towards general protein modulators of FtsZ and identified a combination that anchors filaments to the interface, producing a ring-like phenotype. PUREdrop bridges computational protein engineering and synthetic cell research, elevating the rational engineering of complex biological function to the next level.

## Main

Protein design and optimization promise the generation of proteins with custom properties, opening the door to harnessing biology’s most versatile tools for human engineering ^1,2^. Rapid progress in computational methods has greatly increased success rates and complexity of design ^3–5^, but experimental pipelines to produce and screen must evolve accordingly. Protein-driven bottom-up synthetic biology focuses on finding minimal protein machineries for essential cellular functions, such as division, metabolism, or motility ^6–9^. The field stands to benefit from protein engineering ^1,10^, as the reconstitution of natural proteins have often proven challenging in radically simplified environments ^10,11^. Engineering proteins with bespoke emergent functions, will move the field beyond trial-and-error. Since emergent functions arise from protein self-organization at much larger scales, engineering synthetic cells requires both micrometer-scale spatial control and precise temporal coordination. To screen libraries for complex phenotypes, experimental methods are required that can both controllably generate and quantitatively monitor the emergent dynamics within synthetic cells ^12,13^.

Current experimental methods for screening computationally engineered proteins have focused on molecular functions that can be characterized in bulk, such as minibinders or enzymes with desired catalytic activity ^14–16^. For these applications, semi- and fully automated microtiter-plate workflows have been developed ^17^. Proteins are expressed either in small-volume plates or via cell-free transcription-translation (TXTL) to streamline protein production ^18,19^. This approach provides throughput compatible with computationally designed libraries, while maintaining sequence-to-well identity to readily establish genotype-phenotype relations. However, bulk or ensemble measurements aggregate the entire reaction volume into one averaged signal. This works well for independent single events, such as binders or enzymes, where ensemble averaging is sufficient and the overall readout approximates a Gaussian distribution. In contrast, for higher-order protein functions such as cellular scale self-assembly, these assays would only consider the system in bulk, providing just a single, indiscriminate data point.

Chip-assisted *in vitro* compartmentalization into precisely adjustable droplets helps overcome these limitations by allowing each droplet to act as an individual cell-bioreactor and discrete data point, while also reducing TXTL reagent use ^20,21^. This strategy is mainly applied in ultrahigh-throughput droplet-based screening workflows, where fluorescence-activated droplet sorting (FADS) enables screening of a pooled library containing 10^4^ to 10^6^ protein variants ^22,23^. When coupled with evolutionary approaches that generate library diversity at matching scales, these systems can explore vast sequence spaces ^24,25^. Despite these advantages, such platforms remain limited, relying on a single biochemical signal per droplet at a fixed time point, which in turn prevents the direct mapping of emergent function phenotypes without additional characterization steps after sorting and sequencing ^26^. This exposes a gap in screening methods for emergent function phenotypes (Supplementary Fig. 1), and underscores the need for intermediate approaches that bridge bulk well-plate and droplet-based technologies similar to the MITOMI-based screening approaches ^27–29^, offering the spatiotemporal resolution required, while combining the genotype-phenotype mapping of microtiter plates with the statistical power and efficiency of droplet assays.

To fill the gap and provide comprehensive insights into emergent functions from the onset of expression, we developed PUREdrop, an automated microfluidic platform that produces the designed library within an array of synthetic cell populations organized in a well plate format. PUREdrop sequentially samples distinct protein-encoding DNA from a microtiter plate and co-encapsulates it with a TXTL system into cell-sized water-in-oil droplets. Each droplet population is then allocated to a predefined well downstream. Hundreds to thousands of droplets per variant are then imaged automatically throughout protein expression, allowing parallel observation of emergent microscale functions via time-resolved fluorescence microscopy. As a proof-of-principle, we use the PUREdrop to screen machine-learning re-designed variants of the bacterial cell division protein FtsZ ^30^ for the microscale function of linear self-assembly, comparing the spatial and temporal dynamics to the wildtype. We further demonstrate the PUREdrop use case in screening protein-based modulators of FtsZ and resolving their unique functions by proxy.

### PUREdrop workflow

To enable controlled high-throughput compartmentalization of cell-free expression machinery for diverse gene constructs, we implemented a microfluidic 5-way hydrodynamic focusing design that generates monodisperse water-in-oil droplets. The junction allows the separate and simultaneous introduction of DNA templates (Inner Aqueous 1, IA1) and TXTL reagents (IA2), to produce compositionally distinct synthetic cell populations by exchanging the DNA input (Fig. 1). We used the PUREfrex2.0 system for protein production, due to its defined composition and low background activity. To date, its high-cost relative to the limited number of distinct reactions per kit has represented a major bottleneck to adoption in bulk screening workflows. However, relying on PUREdrop’s droplets-on-demand greatly improves reagent efficiency. The three main PUREfrex2.0 components responsible for TXTL constitute ∼65% of the total reaction volume (IA2), with the remainder comprising DNA and water (IA1). By decoupling the TXTL machinery from the DNA stream, we were able to generate ∼3×10^5^ droplets (∼20 µm diameter) per reaction using only ∼0.87 µL of the PUREfrex2.0 components IA2, in comparison to the minimum ∼6.5 µL TXTL reagents per reaction used in manual emulsification protocols, mounting to 7.4 folds in enhanced reagent efficiency ^13^. This can be further enhanced by decreasing the number of produced picolitre droplets per reaction.

**Fig. 1.**
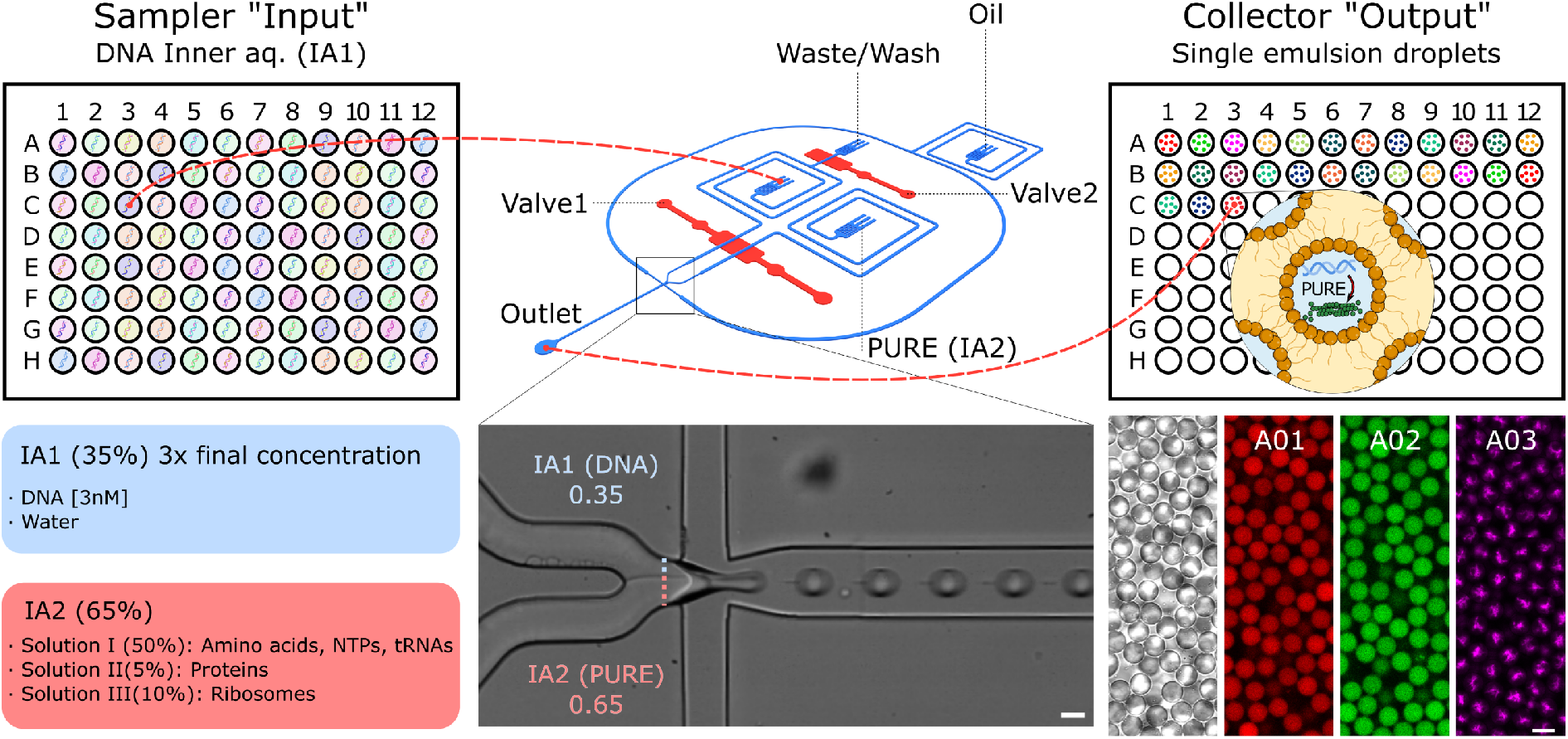
Overview of the PUREdrop microfluidic platform for automated TXTL encapsulation. The schematic illustrates the droplet-based generation of a synthetic cell library, with each well containing a unique construct. A five-way junction produces water-in-oil droplets via pinched flow, mixing IA1 (DNA, autosampler-controlled) with IA2 (PURE TXTL system). The platform integrates a programmable autosampler and fraction collector that multiplex using the well-plate format. A pneumatic valve layer (red) beneath the fluidic main layer (blue) enables stage-specific control via dual control channels. IA1/IA2 compositions are listed. Insets show the microfluidic junction mixing ratio (IA1:IA2 ≈ 1:2), and fluorescence images of droplets expressing sfGFP, mCherry, and FtsZ-Venus. Scale bars represent 20 μm.

To enable autonomous and rapid exchange of DNA constructs in solution IA1 prior to droplet generation, we integrated a 96-well plate autosampler and a fraction collector, placed upstream and downstream of the droplet generation platform, respectively. This setup was adapted from the micrIO system ^31^. We also incorporated a pneumatic valve system with two control configurations, the first mode is for pausing droplet production and diverting fluids off-chip by actuating valve 1 and opening valve 2, while the second mode is for resuming droplet production by reversing the valve actuation. This setup facilitated swift input liquid exchange between samples on chip without manual intervention.

The automated microfluidic workflow integrates three primary operations (droplet production, priming, and cleaning), coordinated through a valve manifold and an electronically actuated fluidic switch. Pressure-driven fluid control, coupled with continuous real-time flow rate monitoring, ensures reproducibility and consistency throughout operational cycles. The fluidic platform is comprised of the microfluidic chip and the external hardware components (Fig. 2a). Figures 2c and 2d present operational data, underscoring the consistency and the duration of representative operational loops between wells, while Fig. 2e provides a detailed breakdown of a single loop.

**Fig. 2.**
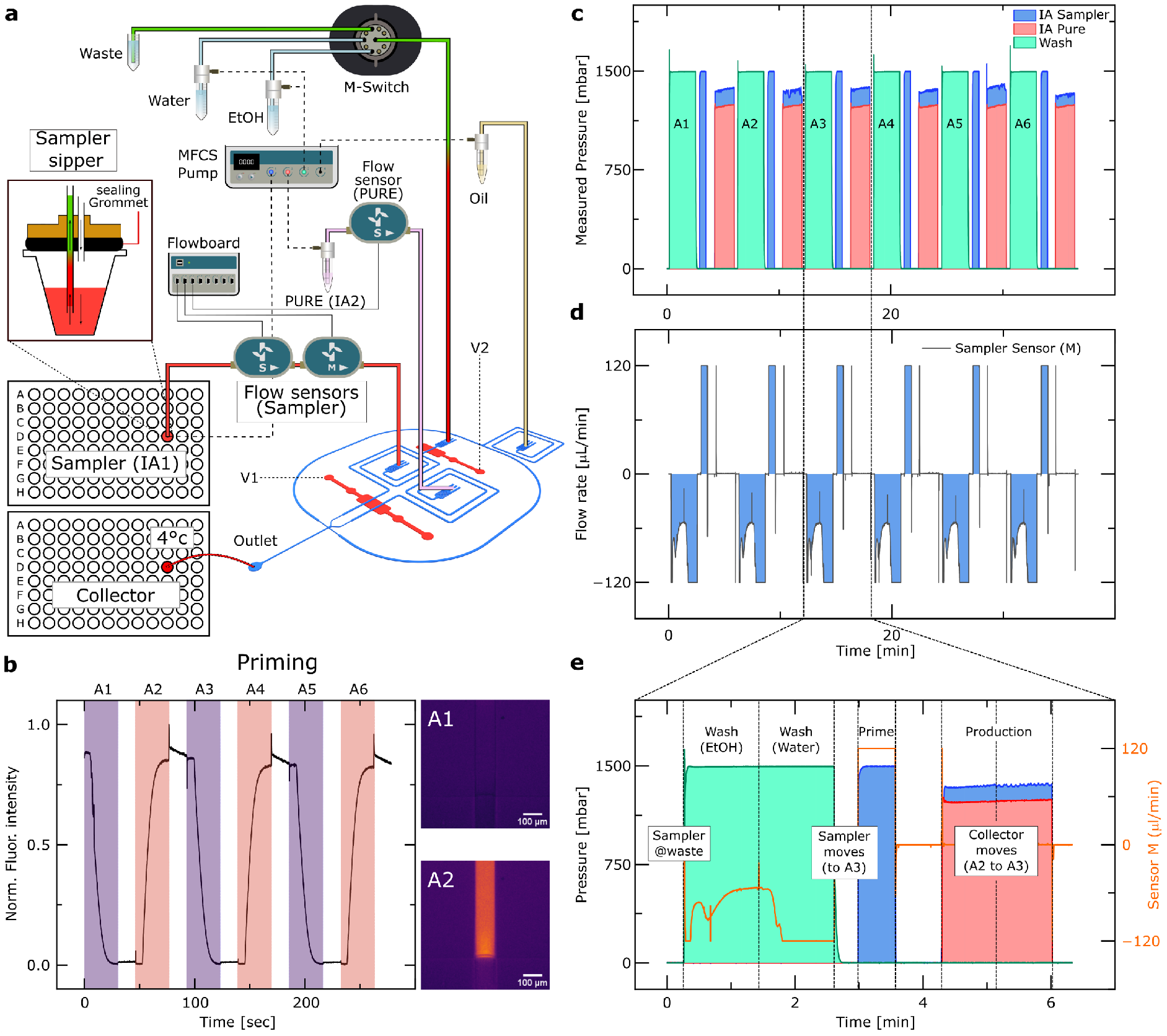
Automated microfluidic platform for synthetic cell production. **a** Schematic representation of the integrated microfluidic platform, comprising the fluidic chip and external hardware components. The platform is driven by pressure-based flow control and coordinated via programmable flow sensors, a valve manifold, and electronically actuated fluidic switches. Sample loading is achieved using an autosampler equipped with a reservoir sealing sipper, which enables positive-pressure extraction from standard 96-well plates without the need for threaded reservoir fittings. Fractions are collected into a cooled (4 °C) destination plate. **b** Priming of the chip with input IA1, monitored by fluorescence readout across wells A1-A6. HPTS dye was loaded into alternating wells (A2, A4, A6) to visualize fluid exchange, with water in intervening wells. **c-e** Real-time monitoring of system dynamics during washing, priming, and production cycles. **c** Measured pressure traces for IA1, IA2, and wash lines. **d** Corresponding flow rate measurements in the tubing segment between the sipper and chip. Positive (+) flow indicates delivery to the chip, while negative (-) flow corresponds to reversed flow directed washing. **e** Detailed temporal profile of a single operational loop (well A3), showing synchronization between pressure and flow sensors. The sipper undergoes sequential ethanol and water rinses prior to sample priming. The fraction collector is actuated mid-cycle to ensure accurate sample deposition.

### Validation of the automated pipeline

To evaluate the reliability and performance of our automated pipeline for droplet-based *in vitro* protein expression, we conducted a validation experiment using DNA-encoded superfolder GFP (sfGFP) and mCherry as fluorescent reporter proteins. These templates were alternately loaded onto 48 wells in the DNA input plate (Fig. 3a). Protein expression was then monitored overnight using automated fluorescence microscopy. As a quality control step, we verified the fidelity of the DNA priming and washing protocols by quantifying the average fluorescence intensity of each imaged droplet across both emission channels in each well at the end of the measurement. Heat maps depicting the per-well average fluorescence intensity showed clear signal segregation between sfGFP- and mCherry-expressing droplets, with minimal cross contamination (Fig. 3b, Supplementary Fig. 2, 3). Representative multi-channel fluorescence images demonstrate the specific protein production from each DNA template (Fig. 3c). A histogram of droplet diameters pooled from 48 wells at the end of the measurement confirmed a unimodal size distribution (20.5 ± 2 µm), consistent with uniform droplet generation and stability (Fig. 3d and Supplementary Fig. 4).

**Fig. 3.**
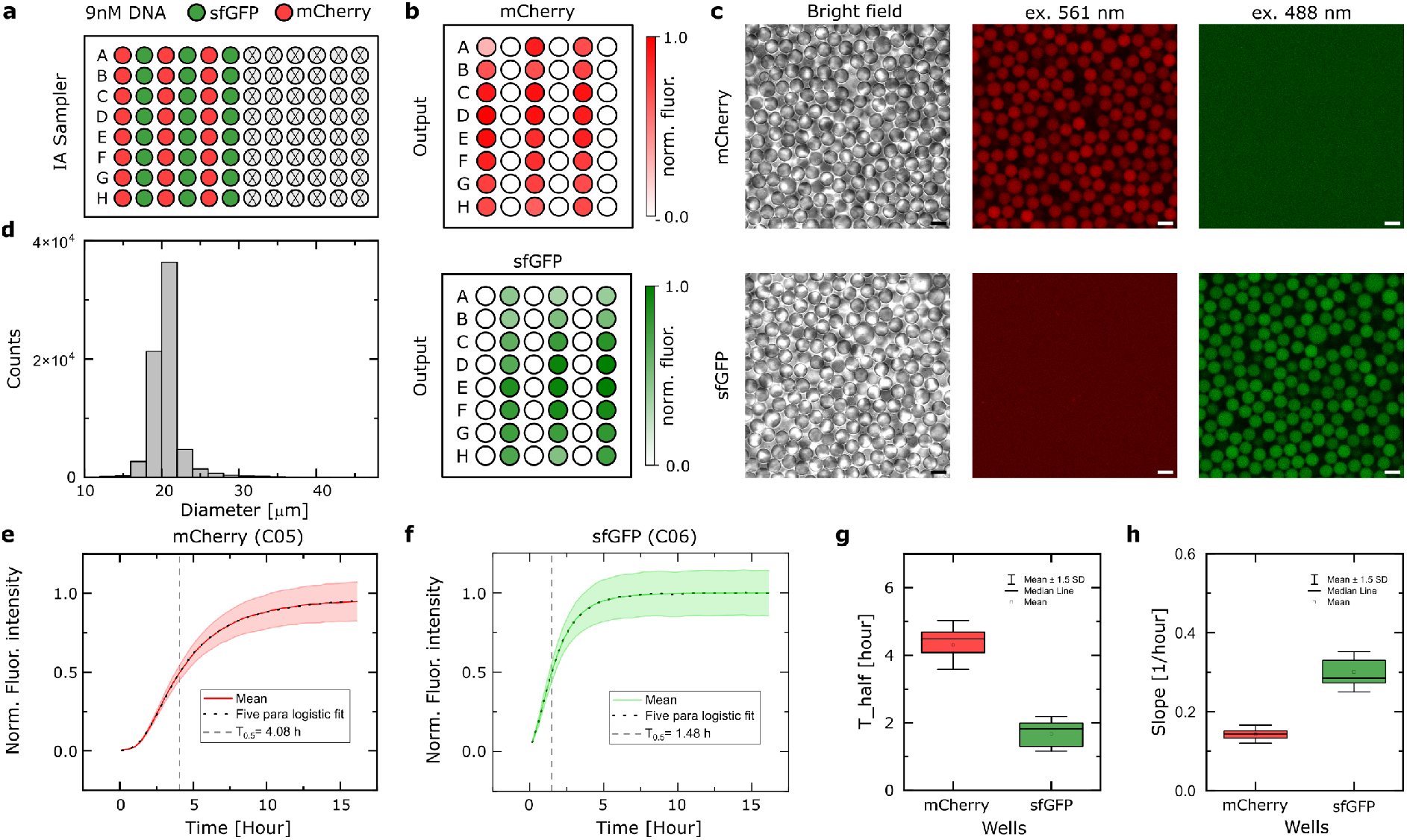
Validation of the automated pipeline using *in situ* protein expression. **a** Schematic layout of the DNA input well plate, showing alternating wells containing sfGFP or mCherry DNA templates (9 nM = 3 nM after encapsulation). **b** Heat maps showing the average fluorescence intensity of all imaged droplets per well, normalized to the minimum and maximum intensity for mCherry and sfGFP, respectively. **c** Representative multi-channel fluorescence microscopy images from alternating mCherry and sfGFP wells. Scale bars, 20 μm. **d** Histogram showing the size distribution of all brightfield imaged droplets, pooled from 48 wells after 15 hours extracted. **e, f** Time-course traces of normalized average fluorescence intensity for droplets in wells C05 (mCherry) and C06 (sfGFP), respectively. Shaded areas represent the standard deviation across droplets C05 (n= 708 ± 12) and C06 (n= 708 ± 11); dotted lines represent five-parameter logistic fits. Dashed lines indicate T_0.5_ values, representing the time to reach 50% of maximum protein expression. **g, h** Box plots comparing T_0.5_ (g) and expression rate at T_0.5_ (slope) (h) between wells expressing sfGFP and mCherry.

Time-lapse image series enabled monitoring of protein expression dynamics within droplets. Normalized fluorescence traces for mCherry and sfGFP wells exhibited asymmetric sigmoidal expression kinetics (Fig. 3e, f), which were captured by five-parameter logistic fits. From these fits, we extracted the time to half-maximal expression (T_0.5_) and expression rate (slope at T_0.5_) as comparative metrics. Across 24 replicates, the extracted T_0.5_ and expression rate slopes were highly reproducible for both fluorophores (Fig. 3g, h). The mean T_0.5_ for synthetic cells expressing sfGFP or mCherry were (1.67 ± 0.34 h) and (4.3 ± 0.48 h), respectively, while the corresponding expression rate were (0.14 ± 0.01) and (0.3 ± 0.03). Expression began consistently across all wells of the same DNA construct following incubation at 30°C, confirming the ability to halt the TXTL machinery at 4°C and trigger it only before imaging. Notably, differences in T_0.5_ and expression rates between sfGFP and mCherry reflect intrinsic differences in folding and maturation kinetics of the two fluorescent proteins ^32^.

### Standardized workflow for screening a construct library

To prepare screening-ready libraries within a few days, we integrated PUREdrop with a quick DNA library assembly protocol (Fig. 4). The process begins with gene fragments containing Golden Gate Assembly (GGA) overhangs, delivered in plates compatible with the ECHO acoustic liquid handler. Using the ECHO, 2 μl GGA reactions containing vector, restriction enzyme, ligase, buffer and selected gene fragments are assembled and incubated at 37 °C for 3 h, after which the products are transformed into chemically competent cells and grown overnight. This part of the workflow builds on and is fully compatible with the semi-automated protein production (SAPP) pipeline ^15^. PURE-compatible DNA templates, comprising a T7 promoter, coding sequence and T7 terminator, are then generated directly from bacterial lysate by PCR amplification. In practice, this allows users following the SAPP workflow to use a small aliquot of the same overnight culture prepared for protein production to also generate templates for PUREdrop, without requiring additional steps. While we implemented a plasmid-based intermediate for robustness and straightforward integration with SAPP, users can also exploit the compatibility of cell-free expression with linear DNA templates to bypass cloning and transformation altogether, enabling an even faster design-to-screen cycle through direct PCR-based library assembly. To obtain sufficient material, multiple PCR reactions were performed in parallel and pooled. The resulting amplicons were purified, quantified, and normalized to 9 nM. Once prepared, a PUREdrop run required 5 hours to encapsulate the contents of 48 wells. The output plate was subsequently imaged through overnight time-lapse microscopy. Using the segmentation software (ArivisPro) it required 1-2 days to extract droplet-level quantitative spatiotemporal datasets and analyze phenotypic behavior.

**Fig. 4.**
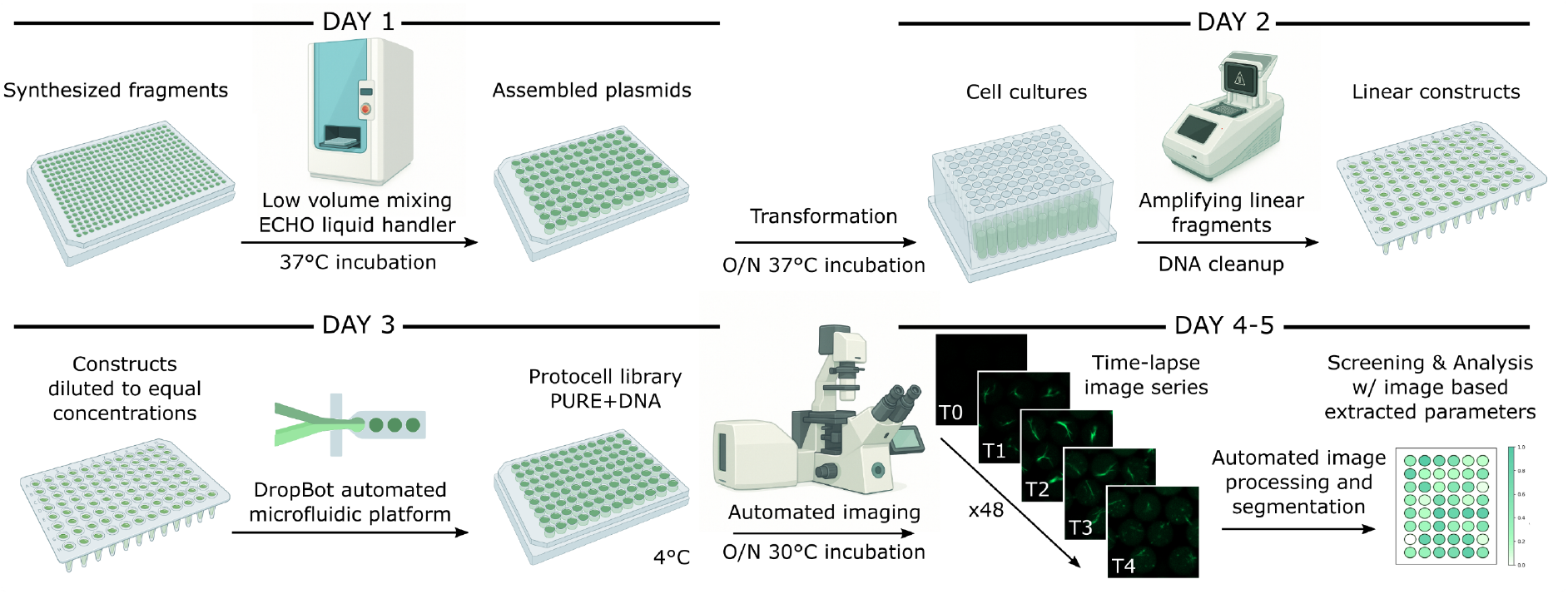
Complete experimental workflow for *in vitro* screening of a construct library. Day 1, Synthesized DNA fragments are assembled into plasmids via Golden Gate assembly using an Echo low-volume liquid handler and transformed into bacterial cells for overnight growth. Day 2, Cells are lysed and linear DNA constructs are prepared by PCR amplification. Crude lysates are filtered to purify the target DNA. Day 3, DNA constructs are normalized to equal concentrations and loaded into the PUREdrop automated microfluidic system. The resulting library is imaged via time-lapse microscopy across 48 wells during overnight incubation at 30 °C. Days 4-5, Time-lapse images are processed and segmented for quantitative screening, enabling construct-level analysis based on image-derived parameters.

### Screening of computationally engineered FtsZ variants

To demonstrate the utility of our pipeline, we applied it to a protein engineering task in the context of building synthetic cell modules, i.e., the *in vitro* re-engineering of FtsZ, a bacterial tubulin homolog involved in cell division^30^. FtsZ is a key cytoskeletal protein that polymerizes in a GTP-dependent manner into dynamic protofilaments. These hundred nanometers long protofilaments can associate laterally and form micrometer-scale bundles ^33^. Our design goal was to preserve the essential functions of FtsZ, including GTPase activity and polymerization capacity, while potentially optimizing expression levels or altering bundling behavior. To this end, we developed a computational workflow to identify mutation-tolerant residues in the FtsZ sequence. Residues were excluded from re-design if they (i) were part of the GTP binding pocket, (ii) took part in the polymerization interface, or (iii) were evolutionarily conserved (Fig. 5a). Depending on the conservation threshold applied, 117 residues (38.2%) or 67 residues (21.9%) were considered safe to mutate. Lower thresholds were avoided, based on previous findings that essential catalytic activity was lost at more permissive cutoffs ^34^. For each conservation level, we used ProteinMPNN ^35^ to re-design the mutable positions. Monomeric structures of all variants were predicted using AlphaFold2 ^36^. Based on mean pLDDT scores, the top 12 sequences from each conservation group were selected for experimental screening (Supplementary Fig. 5), resulting in 24 designed variants (mpnnFtsZv01-24) (Supplementary data. 1). These sequences shared between 73% and 86% sequence identity with wild-type (WT) *E. coli* FtsZ with an average of 79 (50% cutoff) and 47 (70% cutoff) residues being mutated (Supplementary Fig. 6).

**Fig. 5.**
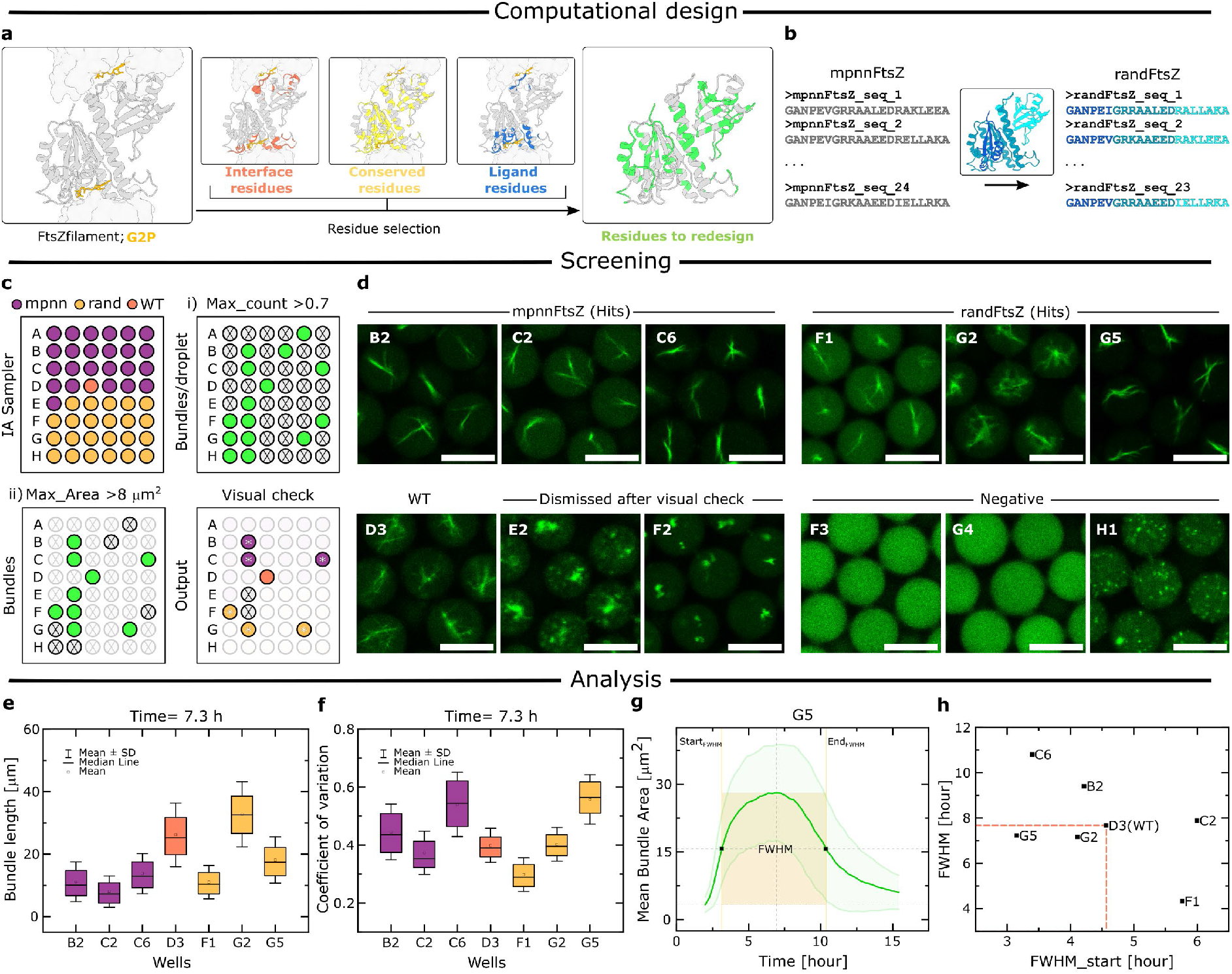
*In vitro* screening of synthetic FtsZ variants for protein bundling activity. **a** Schematic outlining the computational re-design of FtsZ. Residue selection for re-design excluded interface, conserved, and ligand-binding residues, yielding a pool of eligible positions modified using a reverse folding model (ProteinMPNN ^34^). **b** Each of the 24 mpnn-generated sequences was divided into three fragments and recombined in shuffled permutations to generate 23 additional sequences (“rand”). **c** Schematic layout of the DNA input plate, indicating the distribution of wild-type (WT), re-designed, and shuffled re-designed constructs (9 nM). Screening was conducted in three stages: (i) initial selection based on the fraction of droplets exhibiting bundle formation (threshold > 0.7), (ii) exclusion of constructs producing bundles with maximum areas < 8 μm^2^, and (iii) visual curation to eliminate false positives. **d** Representative fluorescence microscopy images from selected wells at 7.3 h, illustrating bundle formation by mpnnFtsZ and randFtsZ hits, WT FtsZ, visually excluded constructs, and negative results. Scale bars represent 20 μm. **e, f** Box plots comparing bundle length and coefficient of variation (CV) of signal intensity for skeletonized bundles per droplet at 7.3 h across all hit constructs. Data were acquired from bundles of WT, B2, C2, C6, F1, G2, and G5, with corresponding N values of 1120, 1206, 1120, 1276, 1013, 1031, and 1264, respectively. **g** Temporal profile of mean bundle area for hit G5. Green shading indicates standard deviation. The yellow region represents the full width at half maximum (FWHM) of the bundle growth and disassembly curve. FWHM_start_ and FWHM_end_ correspond to the time points at which the bundle area first exceeds and later falls below 50% of its maximum value, respectively. **h** Scatter plot mapping FWHM versus FWHM_start_ derived from the temporal profile of mean bundle area for each hit construct. WT FtsZ (D3) is highlighted for reference.

**Fig. 6.**
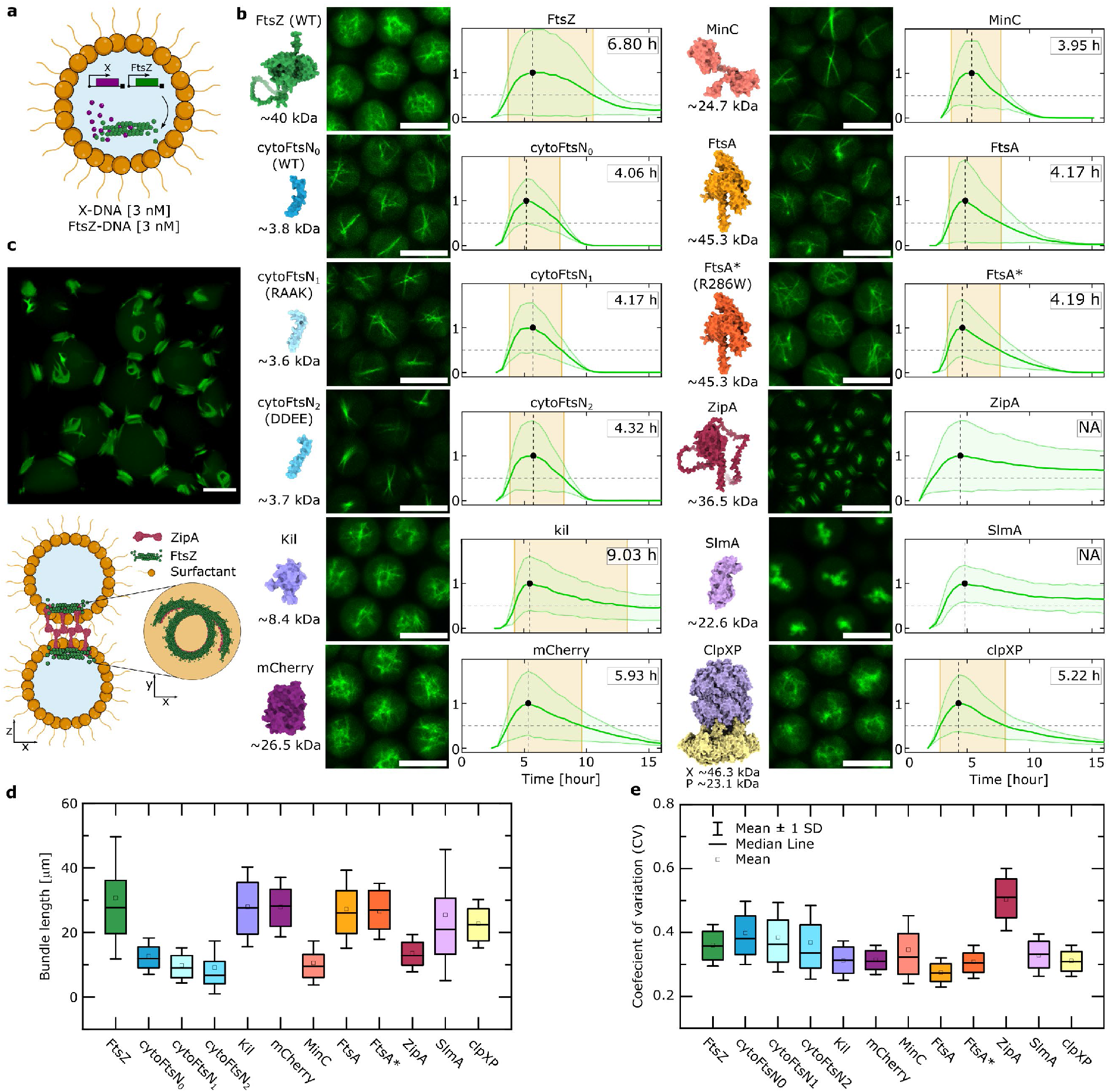
Screening of emergent function modulators. **a** Schematic of co-encapsulation of genes encoding FtsZ and the modulator of interest at similar concentrations (final 3 nM). **b** Structure and size of each protein shown alongside representative fluorescence microscopy images at 5.7 h and corresponding temporal profiles of normalized mean bundle area per droplet. Green shading indicates standard deviation. The yellow region denotes the full width at half maximum (FWHM) of the assembly-disassembly curve. Where applicable, the FWHM value is indicated as the lifetime (h). Scale bars represent 20 μm. Protein structure visualizations were prepared using UCSF ChimeraX 1.11.1 **c** Representative deconvolved z-stack image of a sample expressing ZipA, revealing the three-dimensional arrangement of droplets, with a schematic illustrating localization of ring-like FtsZ bundles at the interface between vertically stacked droplets. **d, e** Box plots comparing bundle length and coefficient of variation (CV) of signal intensity for skeletonized bundles per droplet at 5.7 h across all FtsZ-modulator combinations FtsZ, CytoFtsN_0_, CytoFtsN_1_, CytoFtsN_2_, Kil, mCherry, MinC, FtsA, FtsA*, ZipA, SlmA, and ClpXP, with corresponding N values of 730, 1446, 1323, 1011, 1038, 1156, 1100, 777, 1184, 1011, 813, and 1459, respectively.

All selected designs incorporated the Venus coding sequence between the codons for G55/Q56, allowing visualization, while maintaining high functionality when expressed by TXTL machinery ^9^. For cost-effective and scalable cloning, we split the sequences into three fragments (Supplementary Fig. 7), with junctions strategically positioned at the Venus insertion site and the loop between the N-terminal and C-terminal domains of FtsZ. This fragmentation strategy allowed the synthesis of shorter cost-effective DNA fragments, and the straightforward parallel cloning. All split sites were located in conserved regions, resulting in identical overhangs across all 24 designed sequences. This consistency enabled not only the assembly of each mpnnFtsZ variant, but also the generation of 23 additional chimeric constructs, referred to as randFtsZv01-23 (Fig. 5b), by combinatorially mixing fragments from different designs including WT *E. coli* sequence fragments, resulting in a library of 48 constructs.

To identify FtsZ variants with dynamic bundling behavior, we analyzed the time-lapse fluorescence microscopy data by quantifying the number of segmented fluorescent high-order structures within each droplet coupled with their size, averaged across the field of view at each time point. The screening followed a three-step filtering process (Fig. 5C). First, we selected variants that exhibited high-order structures in over 70% of droplets at any point during the time series, which reduced the initial pool from 48 to 14. Second, we retained only those variants that showed an average structure area exceeding 8 µm^2^ at least once during the time course, narrowing the set further to 8 of 48. Third, manual inspection of the remaining time-lapse sequences was performed and resulted in the removal of two false positives. This yielded six hits in addition to the WT, consisting of three mpnnFtsZ and three randFtsZ variants.

Time-resolved analysis of bundle formation over 15 hours provided further insight into the dynamic behavior of the FtsZ variants from the onset of expression. Bundle area increased steadily, reaching a maximum around ∼7 hours after the onset of expression (Fig. 5f). As a result of GTP depletion, depolymerization overcame polymerization, ultimately resulting in bundle disassembly (Supplementary Movie 1). To compare dynamics across constructs, we quantified the bundle area imaged for each of the hit variants (WT, B2, C2, C6, F1, G2, and G5) (as represented in Fig. 5g). Following the bundle area, we can extract temporal metrics such as full width at half maximum (FWHM), which captures the duration of bundle persistence, and the time to reach half-maximum bundle area, indicating the onset of bundle formation (Fig. 5g and Supplementary Fig. 8). G5 exhibited the earliest onset of bundling (3.15 h), followed by C6 (3.39 h), both faster than WT-FtsZ (4.56 h). Regarding bundle lifetimes, G5 (7.22 h), G2 (7.16 h), and C2 (7.89 h) were comparable to WT (7.67 h), while C6 displayed a longer persistence (10.81 h) (Fig. 5h).

To quantify bundle morphology across the selected hit variants, we analyzed skeletonized bundles at 7.3 h using two complementary metrics. The first is skeleton length, reflecting the spatial extent of the assemblies, and the other is the coefficient of variation (CV) of fluorescence intensity, an image-based proxy for filament bundling ^37^. In this framework, longer skeletons indicate more extended, network-like bundles, whereas higher CV values indicate more compact bundled structures. WT FtsZ (D3) formed long bundles (26.2 ± 10.2), second only to G2, which displayed the most extended structures of all variants (32.8 ± 10.4). G5 also produced relatively long bundles (18.1 ± 7.4), whereas C6, B2 and F1 formed shorter structures (13.8 ± 6.4, 11.2 ± 6.4 and 11.0 ± 5.3, respectively), and C2 showed the shortest bundles overall (8.0 ± 5.0). CV measurements further separated these phenotypes, where G5 exhibited the highest CV (0.56 ± 0.09), followed by C6 (0.54 ± 0.11), indicating more compact and strongly bundled assemblies. B2 also showed elevated bundling (0.45 ± 0.10), whereas WT FtsZ (0.40 ± 0.06) and G2 (0.40 ± 0.06) occupied an intermediate regime, consistent with a more filamentous and less compact organization. C2 showed a lower CV (0.37 ± 0.07), and F1 displayed the lowest CV of all hits (0.30 ± 0.06); together with their shorter structures and delayed polymerization onset, this suggests dynamics shifted away from stable bundle formation and toward filament disassembly. Together, these measurements reveal that the hit variants span a broad morphological space, from extended network-like bundles to shorter, denser assemblies. Notably, the randFtsZ chimeras recovered as hits consistently contained one or two segments derived from either hit mpnnFtsZ variants or the WT sequence (Supplementary Table 1).

These characteristics can be crucial when designing division machinery for synthetic cells and fine-tuning properties such as the thickness of bundles, the timing of bundle initiation, or the persistence of the division Z-ring. The observations provide a valuable foundation for further biophysical characterization of promising variants and for informing parameters in subsequent re-design and screening cycles.

### Screening of emergent function modulators

To demonstrate the versatility of our pipeline, we extended our screening towards additional proteins and peptides that modulate FtsZ self-organization into the Z-ring. Each candidate was tested individually with FtsZ, such that the resulting phenotype reflected the effect of a single co-expressed factor on FtsZ self-organization (Fig. 6a). The readout was based on FtsZ fluorescence, such that changes in bundle formation, lifetime, morphology, and localization served as a proxy for the activity of the co-expressed partner. We selected a compact, functionally diverse first-pass panel of obvious candidates with literature-supported roles in division-site assembly, Z-ring stabilization, antagonism, or turnover ^38^. FtsA and ZipA were included as the two canonical membrane anchors of FtsZ, and FtsA* (R286W) as a gain-of-function comparator ^39–41^. MinC and SlmA represented physiologically distinct inhibitors of Z-ring assembly. MinC weakly antagonizes FtsZ polymerization ^38^, whereas SlmA is a DNA-activated inhibitor that can reorganize FtsZ into condensate-like assemblies in the presence of specific SlmA-binding sequences. CytoFtsN and its RAAK and DDEE variants have been associated with productive proto-ring organization through FtsA, and more recently cytoFtsN has been shown to stabilize and align FtsZ assemblies in reduced reconstituted systems ^42^. ClpXP represented a physiologically relevant regulator of FtsZ turnover ^43^, whereas Kil provided a phage-derived perturbation that disrupts Z-ring assembly *in vivo* in a ZipA-dependent manner ^44^. mCherry served as a non-specific expression control.

In a first comparison of emergent function dynamics, most FtsZ-modulator combinations displayed the familiar assembly-disassembly trajectory from which bundle lifetime could be quantified as the FWHM of the normalized bundle-area curve (Fig. 6b). FtsZ alone showed a lifetime of 6.80 h. Several modulators shifted this profile toward shorter-lived assemblies, most prominently MinC (3.95 h), followed by the cytoFtsN variants FtsN_0_, FtsN_1_ and FtsN_2_ (4.06 - 4.32 h), as well as FtsA and FtsA* (4.17, 4.19 h). ClpXP (5.22 h) and mCherry (5.93 h) had weaker effects, whereas Kil markedly prolonged bundle persistence, producing the longest measurable lifetime (9.03 h). In contrast, SlmA and ZipA did not exhibit detectable disassembly within the imaging window, such that no FWHM could be assigned. These data indicate that the modulators strongly reshape the temporal persistence of the assemblies.

We next quantified bundle morphology at 5.7 h using skeleton length and the CV of the fluorescence signal per droplet (Fig. 6 d,e). In this analysis, longer skeletons are consistent with more extended, branched network-like bundles, whereas shorter skeletons indicate more isolated bundle structures. Likewise, lower CV values reflect weak bundling, whereas higher CV values indicate more compact bundle organization. FtsZ alone formed the longest structures on average (30.7 ± 18.9), consistent with an extended network-like morphology. Several conditions preserved similarly long assemblies, including Kil (28.0 ± 12.3), mCherry (27.9 ± 9.2), FtsA (27.2 ± 12.0), FtsA* (26.6 ± 8.6), SlmA (25.4 ± 20.3) and clpXP (22.8 ± 7.5), whereas the cytoFtsN variants and MinC shifted the population toward shorter structures, with mean lengths of 9.2-12.7 for FtsN0/1/2 and 10.6 ± 6.8 for MinC. These shorter lengths are consistent with reduced higher-order network formation and aligned bundles.

The CV values further separated these phenotypes. FtsA showed the lowest mean CV (0.27 ± 0.05), followed by FtsA* (0.31 ± 0.05), clpXP (0.31 ± 0.05), Kil (0.31 ± 0.06) and mCherry (0.31 ± 0.05), indicating comparatively weakly compacted, more filament-like assemblies. FtsZ alone occupied an intermediate regime (0.36 ± 0.07), similar to MinC (0.35 ± 0.11), SlmA (0.33 ± 0.07) and the cytoFtsN variants (0.37-0.40). In contrast, ZipA was clearly separated from all other conditions, displaying by far the highest CV (0.50 ± 0.10) together with relatively short bundle lengths (13.6 ± 5.8). Most notably, ZipA emerged as a standout hit of the screen. Beyond forming relatively short, highly compact bundles with no detectable disassembly, ZipA anchored FtsZ assemblies specifically at the droplet-droplet interfaces from the onset of expression. Despite the fact that these synthetic compartments were stabilized only by surfactant, ZipA nonetheless displayed transmembrane-like localization behavior, partitioning selectively to interfacial regions. At the interface between vertically stacked droplets, this spatial configuration enabled clear visualization of a ring-like bundle morphology (Fig. 6c). These results illustrate the diversity of phenotypes that can be observed with the PUREdrop and underscore the effectiveness of our cell-free imaging strategy in tracking protein higher-order emergent function.

## Conclusion

Machine learning-aided design of emergent protein functions for deployment in well-controlled synthetic cell environments represents a central frontier in bottom-up synthetic biology, with implications spanning fundamental biological insight to powerful cell-free applications ^29^. A great bottleneck of this new research direction is the availability of standardized and comprehensive screening routines that provide direct access to spatiotemporal readouts. To serve the needs of this challenge, we developed the PUREdrop platform, an automated microfluidic routine for generating and organizing synthetic cell populations in a well plate format. We described how to integrate parallelized cloning, DNA amplification, PUREdrop and imaging into a streamlined workflow capable of generating high-quality, time-resolved data on protein functions that dynamically unfold within 24 hours. Co-encapsulating protein-encoding DNA with the TXTL machinery not only offers a standardized, efficient route to produce the protein of interest, but also acts as a temperature control to enable the parallel screening of sequentially prepared libraries and capture their higher-order functions throughout protein expression.

For a first showcase study, we used the platform to screen a library of 48 computationally re-designed variants of the filament-forming protein FtsZ, a central protein used in bottom-up synthetic biology for creating a minimal cytoskeleton-based cell division machinery ^8,9^. We identify and characterize 6 variants, which show altered expression dynamics, assembly kinetics and bundling phenotypes compared to the wildtype. Four variants showed earlier onset of bundle formation, and two variants assembled into larger bundles. Beyond our interest in engineering physical characteristics of these bundles, their dynamics can have substantial implications on the cellular resources allocated to give rise to the desired bundles. This becomes critical when integrating FtsZ with other protein systems within a single compartment, a major challenge for building a multi-functional synthetic cell ^10,45^. Beside designed libraries, the focus of screening within TXTL-based synthetic cells, can be shifted towards optimizing protein expression systems ^46^, testing environmental conditions or systematically explore concentration ratios of multiple DNA templates for co-expression, and deduce how the metabolic burden shifts when multiple variants are introduced. We explore this latter application in our second showcase study. Here, we applied PUREdrop to screen a functionally diverse panel of proteins and peptides that modulate the FtsZ-ring assembly, extending the platform from single-protein library screening to the analysis of emergent behavior in multi-component synthetic cells. By co-expressing each candidate individually with fluorescently labelled FtsZ, we quantified its impact through the resulting FtsZ phenotype, including changes in bundle morphology, persistence and spatial organization. The screen captured a wide spectrum of behaviours, ranging from accelerated disassembly to highly persistent, compact assemblies. Of all candidates, ZipA stood out by promoting interface-anchoring, ring-like FtsZ bundles despite the surfactant-stabilized nature of the compartments. Together, these findings demonstrate that PUREdrop can be used not only to screen for optimized protein variants, but also to uncover productive combinations of interacting factors that drive higher-order cytoskeletal organization in synthetic cells.

A major advantage of PUREdrop’s on-chip encapsulation is its substantial reduction in reagents used per variant. We imaged only a few thousand droplets out of ∼300k generated, corresponding to an estimated 7.4-fold reduction in PURE reagent consumption compared to conventional manual protocols. By implementing a droplets-on-demand modality and conserving TXTL reagents throughout operation, total reagent consumption scales directly with the number of picolitre droplets produced for each population, offering substantial flexibility to balance throughput, data resolution, and resources.

Looking ahead, we anticipate a range of developments and applications including the PUREdrop pipeline. In synthetic biology, advancing from single emulsion synthetic cells to double-emulsion templated Giant Unilamellar Vesicles (GUVs) will be a key next step ^47,48^. This transition would allow screening to be carried out in more cell-mimicking, membrane-bound environments. On the computational side, PUREdrop could be coupled to active-learning workflows such as METIS and EvolvePro ^49,50^, enabling iterative cycles in which quantitative data guide the selection of the most informative parameters for subsequent rounds. In therapeutics discovery, where bundles and higher-order protein assemblies serve as targets for antimicrobials ^51,52^, the pipeline could support the characterization of designed protein or peptide libraries to identify and study *de novo* inhibitors or modulators of essential emergent function.

## Methods

### PUREdrop automation routine

#### Droplet production

Pressure-driven droplet production was regulated using the two inline flow sensors (S) positioned to control both the IA1 and IA2, which provided real-time feedback for automated pressure adjustments. This allowed for maintaining a droplet (approx. ⌀20 μm) generation rate of ∼3 kHz, while constraining the flow rate ratio between the two at ∼1:2 (Supplementary Movie 2). Through the sensors we defined a combined volumetric flow between the inner aqueous solutions (IA1 and IA2) of 0.8 μL min^−1^. Owing to laminar flow conditions within the microchannels, component mixing occurred only after droplet formation (as imaged in Fig. 1). Droplet production runs lasted 100 s per well, strategically timed to ensure the output tubing was never fully filled with droplets, thereby avoiding cross-contamination between wells. The fraction collector was programmed to reposition the target well midway through the droplet production interval. The collected droplets are then maintained at 4 °C to halt TXTL activity during the automated run. We aimed at the production of ∼300K droplets per well to ensure full coverage within the observation well regardless of the imaged area (Supplementary Fig. 9).

To maintain droplet stability over 15 hours during incubation at 30°C and to enable direct imaging in standard 96-well plates, we could not rely on commercially available fluorinated oil-surfactant combinations (density >1.6 g/cm^3^), as such droplets tend to float, impeding straightforward imaging. Although imaging of floating droplets is technically feasible, it requires manual handling to transfer to cell counting slides or capillaries, rendering screening workflows cumbersome and inefficient for large datasets. To circumvent this, we adapted the continuous oil phase composition following previous protocols ^53^, using mineral oil (density 0.88 g/cm^3^) paired with Span 80 and Tween 80 surfactants (4.5% and 1.5%, respectively). This formulation provided stable droplet morphology at 30°C without observing major coalescence events and minimal shrinkage across the droplet interface during the observation time (Supplementary Fig. 4).

#### Priming

The priming process entails loading the chip with the target IA1 solution from the input plate. This is achieved by applying the first valve mode that permits fluid flow through and off the chip until the incoming IA1 solution from the autosampler carrying the desired DNA completely displaces any residual fluid. The discarded solution is collected in the waste reservoir post the M-switch. The fluidic path carrying IA1 from the input well to the chip has an estimated dead volume of 42 µL, and the system can sustain a maximum operational pressure of 1500 mbar against the sealing sipper without air leakage. The time required for complete solution replacement was determined experimentally with fluorescence imaging (Fig. 2b). This process involved alternating the chip priming between wells containing HPTS dye (even-numbered) and water (odd-numbered). Fluorescence intensity shifts were observed within 30 s, and we selected a conservative priming duration of 35 s to ensure reliable and reproducible fluid replacement.

#### Cleaning

A cleaning protocol was implemented to minimize the risk of DNA cross-contamination between input wells. Sequential washes with 50% ethanol followed by water were performed, with each step lasting 70 seconds to clean the fluidic path from the sipper to the chip. This was achieved by applying 1500 mbar pressure to both reservoirs, while the M-Switch dynamically selected the appropriate fluid source during each cycle. This corresponds to dispensing 97 µL of ethanol solution followed by 162 µL of water through the priming path per washing cycle.

The M-switch configuration supports up to nine distinct cleaning solutions, enabling flexible and programmable cleaning protocols tailored to specific experimental requirements. Fig. 2d presents real-time flow rate data from the Sampler sensor M, illustrating the priming and washing phases. Positive flow rates indicate forward delivery during priming, while negative values correspond to reverse flow during washing, as cleaning solutions are directed toward the waste reservoir positioned adjacent to the IA sampler input plate. The transition from aqueous to ethanol can be observed by a shift in measured flow rate from 120 µL/min to ∼60 µL/min, with the reverse indicating the removal of the ethanol. Although the sensor is capped at ±120 µL/min and calibrated primarily for aqueous solutions, limiting its quantitative accuracy for ethanol, the observed change reliably confirms successful fluid exchange between the cleaning liquids.

The operational loop requires 375 seconds per well and begins with the washing step (140 s), followed by priming (35 s) and then droplet production (100 s). Throughout all three stages, oil is continuously perfused through the chip serving multiple important functions. In addition to its essential role in droplet generation, the oil clears residual droplets from the output tubing and forms a stabilizing barrier that prevents evaporation during overnight measurements. In the present study, although droplets were sorted into a 96-well plate format, only 48 wells were populated per run to balance preparation time with downstream time-lapse imaging and temporal resolution. There is considerable room for optimization to reduce the total runtime and scale up to 96-well preparation within the same timeframe. For instance, increasing the flow rates during the washing steps, refining mechanical movements (58 s) and reducing the production time could cut those times in half, ultimately enabling faster and larger-scale library preparation.

### Microfluidics setup and operation

The oil phase was prepared by mixing 4.5% (v/v) Span-80 (Sigma-Aldrich) and 1.5% (v/v) Tween-80 (MP Biomedicals) in mineral oil (HP50.3, Carl Roth GmbH, Germany). Homogeneity of the mixture was maintained by continuous stirring using a magnetic wing stirrer bar (16 mm × 10 mm, VWR, UK) prior to and during its introduction into the microfluidic chip. Cell-free protein expression was conducted using the PUREfrex2.0 system (GeneFrontier, Japan), consisting of two distinct aqueous solutions, designated as inner aqueous 1 (IA1) and inner aqueous 2 (IA2). IA1 contained linear DNA constructs diluted to a final concentration of 9 nM in nuclease-free water (DNA preparation described in the SI). Aliquots of 100 μL per well were dispensed into a PCR input plate (ThermoScientific, AB-0800) and subsequently sealed with aluminum foil (ThermoScientific, AB-0626). IA2 was prepared by premixing PURE solutions I, II, and III, and maintained at ∼ 4 °C throughout the experiments using a Thermomixer C (Eppendorf) equipped with a custom-drilled lid. The dead volume allocated within tubing and connections, can be minimized by using smaller inner diameter tubing. All fluid reservoirs, excluding the input plate, were pressurized using Fluigent Fluiwell fittings. The microfluidic chip (fabricated as described in the SI) was mounted directly on a microscope stage. Valve control dead-end channels were primed with water, which was introduced by pressurizing fluid reservoirs (Microtube, PP, 0.5 mL, Brand) connected to the chip via polymer tubing (Masterflex Tygon tubing, 0.02” × 0.06”, Fischer) fitted with right-angled blunt steel tips (Darwin Microfluidics, UK). Pressure lines were regulated using a P-Switch, requiring a dedicated MFCS channel.

Two wash reservoirs containing either 50% (v/v) ethanol or water were pressurized from a single MFCS channel, which was split into two separate pressure lines, each connecting via Tygon tubing to an M-switch. The main fluidic line was the M-switch was connected to the chip wash inlet through PEEK tubing (510 µm × 255 µm, IDEX). Waste fluid from the priming process was directed into a dedicated reservoir through the M-switch, while waste generated during washing was collected in a reservoir mounted on the autosampler stage. The oil, IA1, and IA2 reservoirs were independently pressurized through dedicated MFCS-EZ system channels. IA1 was delivered from the input plate to the chip through PEEK tubing equipped with two inline flow sensors (one Flow Unit S and one Flow Unit M), whereas IA2 utilized a single inline flow sensor (Flow Unit S).

The output plate utilized was a Sensoplate microplate (96-well, flat-bottom glass, Greiner Bio-One) maintained at ∼4 °C using a ColdPlate thermoblock (QINSTRUMENTS GmbH, Germany). The plate was covered by a custom-built lid providing continuous nitrogen flow to prevent water condensation, featuring an opening for the stationary fraction collector arm. The output plate assembly, including the ColdPlate and custom lid, was securely mounted onto the moving XY-stage of the fraction collector. Spatial layouts of both input and output plates, critical for precise XY-stage alignment, were defined and imported via plate maps.

### Automation assembly

The autosampler and fraction collector were assembled as described previously ^31^, except where modifications are specified here. Both sampler and collector systems incorporated a motorized XY-stage (Scan IM 120 × 100, Marzhauser GmbH) controlled by stage controllers (TANGO 2 DT). For the autosampler, a single-axis Z-stage (LIMES 150-300-HiSM, OWIS GmbH, Germany) was added, connected to a PS 10-32 controller and mounted onto a Z-stage bracket (MONT-LIMES150-Z). All stage controllers were connected to a PC through USB-serial adapters. The motorized stages were positioned on both sides of an inverted microscope (Olympus IX71, 20× objective, FASTCAM Mini AX200) and mounted on Ø1″ pillar posts, aligning them vertically with the microfluidic chip on the microscope stage (Supplementary Fig. 10). The sample sipper and collector tubing holder were machined or 3D-printed from STL models available in the micrIO GitHub repository^31^. The sipper assembly comprised a rubber grommet (1/4″ I.D.) and two metal connector tips machined from 23G needles (B. Braun SE), each blunt-ended and 40 mm in length. These tips were secured onto and through the PEEK sipper body using cyanoacrylate adhesive. One tip, used for liquid aspiration, was positioned to pierce through the foil seal of each well and reach its bottom. The second tip, supplying pneumatic pressure, was bent at a 90° angle and positioned above the foil surface. A 3D-printed sampler arm attached to the Z-stage enabled precise vertical movement, allowing the sipper to seal each target well during sampling and retract between sampling steps or move to a waste position. Fluid handling was driven by MFCS-EZ pressure pumps (Fluigent S.A., France) operated in conjunction with a microfluidic bidirectional valve (M-Switch), three flow sensors (two Flow Unit S and one Flow Unit M) addressable through a sensor hub (Flowboard), and a pneumatic valve controller (P-Switch). The entire fluidic system was controlled either through the OxyGEN software interface or programmed using the Fluigent SDK. Automation routines for both the autosampler and fraction collector were implemented using a customized version of the open-source Python package acqpack ^31^, updated for compatibility with Python 3.11.5. This package coordinates stage movements and microfluidic operations by incapsulating hardware commands into user-friendly high-level functions. It also enables the loading of hardware configuration files containing instrument parameters and experimental settings. All stages of the experimental workflow, from protocol design through execution, were managed within a Jupyter notebook environment. The corresponding code is available in the GitHub repository: https://github.com/KANahas/PUREdrop.git.

### Imaging, image segmentation and analysis

Automated imaging of the output plate was conducted overnight (∼15 hours) using a Zeiss LSM980 confocal laser scanning microscope equipped with a Plan-Apochromat 20x/0.80 air objective M27 (Carl Zeiss AG, Germany). Fluorophores were excited using a 488 nm laser for sfGFP and Venus-FtsZ constructs, and a 561 nm laser for mCherry. Imaging intervals for each well were 25.3 minutes for the dual fluorescence imaging experiments and 17.5 minutes for the FtsZ variants screen. In parallel, brightfield images were acquired using the microscope’s transmitted photomultiplier tube (T-PMT). The plate was mapped out using the AI sample finder tool and positions of ROIs were recorded. Each well was imaged across 2 tiles for the sfGFP-mCherry validation and 4 tiles for the FtsZ variants experiments, with each tile covering an area of 400 × 400 µm and a spatial resolution of 0.15 pixels per µm. The microscope incubation chamber was controlled at 30 °C and focus strategy utilized Definite focus 3.0 to maintain focus throughout the imaging process.

Image analysis was performed using ArivisPro 4.3. Image channels were first selected using the selection module to isolate GFP and brightfield channels. Segmentation was performed in two stages. First, droplet masks were generated from the brightfield channel using Cellpose with the pretrained “cyto2” model. Subsequently, a machine learning-based segmentation of the GFP channel was performed using the Ilastik module, where a trained classifier distinguished protein bundles from the background to generate initial object masks. Following segmentation, feature-based filtering was applied to exclude irrelevant objects based on size and shape, including removal of those touching image edges via the touching edge filter. A second round of morphological refinement was performed, including smoothing and hole-filling operations on the protein bundle masks. Protein bundle objects with a projected area of less than 0.18 µm^2^ were excluded from downstream analysis. To determine spatial relationships, the compartmentalization module was used to associate internal protein bundles with their corresponding droplet masks, enabling hierarchical tracking of bundle counts and identities within individual droplets. For skeleton extraction at a single time point, a second segmentation using an intensity threshold-based workflow was implemented. The GFP channel was first denoised using a discrete Gaussian filter, followed by filament-enhancing shape detection. Segmentation was then performed using automatic intensity thresholding with the Li thresholding method to generate filament masks. Skeletons were extracted from the segmented filament masks using the Segment Morphology module with medial-axis skeletonization performed plane-wise for the single analyzed time point. The resulting skeleton objects were then associated with their parent droplet and filament objects for downstream analysis. Quantitative features such as object area, mean fluorescence intensity, and skeleton-associated measurements were exported and saved as CSV files. This pipeline was applied to the full dataset, encompassing 48 tile scans across multiple time points. Processed image data were analyzed using a custom Python script to organize, filter, and pivot object-level information for downstream statistical analysis.

### Computational Design of mpnnFtsZ

The computational FtsZ optimization pipeline was inspired by Sumida *et al*. ^34^. In short, FtsZ was analyzed for (i) conserved residues, (ii) residues taking part in ligand binding, and (iii) residues taking part in the polymerization interface. A cryo-EM structure of a FtsZ filament (PDB: 8IBN) was then provided as input to ProteinMPNN ^35^, where the residues previously selected were fixed, and all other residues were re-designed.

#### Sequence conservation analysis

To find residues that are evolutionary conserved, we generated a Multiple Sequence Alignment (MSA) of sequences that are at least 30% identical to the *E. coli* FtsZ sequence and calculated the entropy per column of the MSA as measure of conservation. All sequences stored in the UniProtKB database ^54^ with gene name “FtsZ” were downloaded (download date: 2024/12/20). Sequences were clustered with mmseqs2 ^55^ with identity threshold 0.3, and only sequences in the same cluster as the *E. coli* FtsZ (UniProt ID P0A9A6) were kept. These sequences were again clustered by mmseq2, with identity threshold 1.0, to avoid duplicates. The MSA was generated using MAFFT ^56^, run with flags “--localpair --maxiterate 1000”. For each position in the MSA, the entropy was calculated with a custom python script implementing

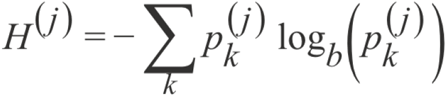

where *H*^*(j)*^ is the entropy for column *j, p*_*k*_^*(j)*^ is the frequency of amino acid *k* in column *j*, and *b* is the base (here *b*=21 for amino acids). The final conservation score C^(j)^ for column *j* is then calculated as

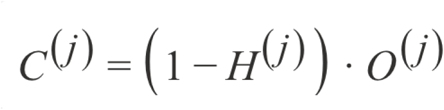

where *O*^*(j)*^ is the fraction of non-gap characters in column *j*:

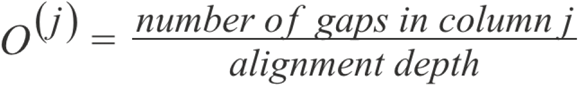

This makes columns that are mostly gaps less conserved (Supplementary Fig. 11). To select conserved residues, the 50% or 70% most conserved residues of the *E. coli* FtsZ sequence were selected. For this selection, the first 10 residues and the last 67 residues of the *E. coli* sequence were ignored, as they were not part of the pdb file used as input to ProteinMPNN.

#### Selecting ligand & polymerization interfaces

The structure of the FtsZ filament from *Klebsiella pneumoniae* (PDB ID: 8IBN), which has 99% sequence identity to *Escherichia coli* FtsZ in the solved parts, was analyzed using a custom python script (Supplementary Fig. 12). A 7 Angstrom radius sphere was specified centered around each atom of the GTP binding pocket, and all residues within at least one of these spheres were considered ligand binding residues and fixed. Similarly, for the polymerization interface, a 7 Angstrom radius sphere was specified centered around each atom of each chain, and residues of other chains having at least one atom within at least one of these spheres were considered polymerization interface residues and fixed.

#### Running ProteinMPNN

The indices of the residues selected to be fixed by conservation, ligand interaction, and polymerization interface were saved as comma-separated list. Additionally, indices 51, 52, 53, 54, 55 and 56 were also fixed, to allow standardized cloning to introduce Venus (3 amino acids before and 3 amino acids after the planned Venus insertion, such that standardized overhangs could be used) (Supplementary Fig. 11). ProteinMPNN ^35^ was run using chain B from PDB 8IBN as input, fixing selected residues, excluding amino acids M and C, sampling from temperatures 0.1, 0.2, and 0.3, using batch size 1 and seed 42, and generating 16 sequences per parameter set. This resulted in 96 sequences (3 temperatures * 2 conservation cutoffs * 16 = 96).

#### Selection of Sequences to order

A local install of ColabFold ^57^ was used to predict the structure of the optimized FtsZ variants. Importantly, the first 11 and last 67 residues of the full *E. coli* sequence were missing from the generated sequences (as described above). These residues are highly unstructured and thus badly predicted by AlphaFold. As this would mostly introduce noise, we ignored it for scoring, and did not include these residues in the structure prediction. Predictions were split into two batches based on conservation threshold, and the 12 sequences with highest average pLDDT score ^36^ per batch were selected for ordering (Supplementary Fig. 5).

#### mpnnFtsZ DNA sequence ordering

The missing first 11 and last 67 residues of the wildtype *E. coli* sequence were appended to the selected sequences. The resulting 48 amino acid sequences were reverse translated and codon optimized with a custom python script. Each sequence was split into three fragments, always at the same index. This split index was selected such that the flanking at least four bases were identical for all constructs, and the desired fragments from different sequences could be assembled using Golden Gate cloning ^58^ (Supplementary Fig. 7). Golden Gate restriction sites plus short random sequences were added to the fragments, and the gene fragments were ordered from GenScript as Titan Gene Fragments in a 384 well plate in TE Buffer low EDTA (10 mM Tris-Cl pH 8.0, 0.1 mM EDTA; 20 ng/μl, 500ng yield). The Venus fragment was ordered as gBlocks HiFi from IDT and diluted to 40ng/μl in nuclease-free H2O.

## Data availability

The source data generated in this study are provided in this paper. The original microscopy timelapses are available under restricted access due to their large file size (> 200 GB). These data are freely available by contacting the corresponding author. The corresponding code is available in the GitHub repository: https://github.com/KANahas/PUREdrop.git.

## Acknowledgements

Funding was provided by the European Research Council (ERC Synergy Grant MetaDivide, no. 101167181). BPF is part of the Graduate School of Quantitative and Molecular Biosciences Munich (QMB). We would like to thank MPIB Imaging Facility (RRID:SCR_025739) for help with the image analysis workflow. We thank Lukas Milles for help with and advice on the SAPP protocol, providing the target vector and training on the ECHO liquid handler. We thank Michaela Schaper for help with cloning and sample preparation.

