## Supplementary information for "Automated Synthetic Cell-based Screening for Designed Proteins with Emergent Functions"

### **Supplementary information:** Automated Synthetic Cell-based Screening for Designed Proteins with Complex Functions

[1]Dept. Cellular and Molecular Biophysics, Max Planck Institute of Biochemistry; Martinsried, D-82152, Germany

*Corresponding authors

#### Mold fabrication

The chip is a multilayered polydimethylsiloxane (PDMS) device fabricated using standard soft-lithography techniques. Two molds were produced: one corresponding to the main chip (top layer) and the other to the valve control layer (bottom layer). The main chip channels were formed using a master mold with two distinct channel profiles: (1) rectangular features with heights of ~37 µm, and (2) rounded features with heights of ~53 µm (Supplementary Fig. 13). The mold was fabricated using a two-photon polymerization printer (Photonic Professional GT2, Nanoscribe GmbH) and IP-Q resist, which enabled the creation of both rectangular and rounded channels in a single step from a 3D STL AutoCAD design. To enhance structure adhesion to the substrate a 4″ silicon wafer (University Wafer, USA) was Oxygen plasma treated (5 min at 0.3 mbar, 50% power; ZEPTO, Diener Electronic, Germany) and subsequently placed in 3-(Trimethoxysilyl) propyl methacrylate dissolved in toluene for 1 h (Gernhardt et al., 2020). The printed substrate was developed in PGMEA and subsequently silanized with 1H,1H,2H,2H-perfluorooctyltrichlorosilane (Thermo Scientific). The rectangular features for the valve control layer have a height of ~35 µm and were realized by spin-coating SU-8 3050 negative photoresist (MicroChem Corp.) at 3000 rpm for 60 s with a ramp of 100 rpm $s^{-1}$ on a 4″ silicon wafer. Spin coating was followed by a soft bake (1 min at 65 °C, 5 min at 95 °C). The resist-coated substrate was exposed using a maskless laser writer (µPG101, Heidelberg Instruments), followed by a post-exposure bake (1 min at 65 °C, 5 min at 95 °C). The structure was developed in PGMEA and hard baked (30 min at 140 °C). Heights and profiles of the molds were measured using laser profilometry (VK-X1100, Keyence, Japan; Supplementary Fig. 13).

#### Preparation of replica PU molds from PDMS chips:

Polyurethane (PU) molds were prepared using cured PDMS chips as master templates. Smooth-Cast™ 310 (Smooth-On Inc.) polyurethane resin was prepared by mixing **30 g of Part A with 27 g of Part B** according to the manufacturer's recommended ratio. The mixture was prepared using a planetary vacuum mixer (ARV-310, Thinky Corp., Japan) to remove entrapped air before being slowly poured over the PDMS master. The resin was allowed to cure at room temperature until fully hardened. Once cured, the PU mold was gently peeled from the PDMS chip, yielding a rigid negative replica of the fluidic channels. (Desai et al., 2009)

#### Device fabrication:

PDMS replicas of the main chip were obtained by mixing the elastomer with curing agent (Sylgard 184, DowSil) in a 9:1 ratio, homogenized and degassed simultaneously for 2 min using a planetary vacuum mixer (ARV-310, Thinky Corp., Japan). The mixture was poured onto replica molds and baked for at least 2 h at 75 °C. The valve control PDMS membrane was prepared by spin-coating PDMS at 2000 rpm for 30 s (ramp 500 rpm $s^{-1}$). After curing and peeling off the main chip, inlets and outlets were punched using a 0.5 mm biopsy punch tip (WPI, UK) adapted to a home-built drill puncher. A glass slide (76 mm × 26 mm) was coated with PDMS and used as the base. The main chip was aligned and bonded to the valve control membrane by exposing both surfaces to oxygen plasma (10 s at 0.3 mbar, 50% power), followed by thermal annealing at 75 °C for 10 min. After bonding, the main chip was peeled off and valve control inlets were punched. The assembly was then bonded to the PDMS-coated glass slide via a second oxygen plasma treatment step.

**Z-stack deconvolution:**

Stack images acquired with the Zeiss Airyscan 2 microscope were deconvolved with Huygens Essential version 25.10 using the "Standard" Deconvolution Express strategy (Scientific Volume Imaging, The Netherlands, http://svi.nl).

**DNA preparation:**

The cloning procedure is inspired by the Semi Automated Protein Production (SAPP) protocol (Qian et al., 2025). On Day 1, the linear fragments delivered by GenScript (fragments 1, 3 and 4) and IDT (fragment 2) were mixed with a Golden Gate Assembly (GGA) Master Mix in an Armadillo 96 well plate using an Echo 525 Acoustic Liquid Handler (Beckman Coulter). The target vector was LM627 (Addgene), stock concentration 666 ng/µl, which contains a C-terminal SNAC tag followed by a 6xHis tag. As fragment 4 was ordered with a stop codon, these tags were not expressed. Composition of a single GGA reaction is depicted in Supplementary Table S2. After mixing, the Armadillo plate containing the 48 reactions was covered with a PCR plate seal and incubated at 37°C for 4 h. Then, the assembly reactions were directly used to transform 12 µl of NEB 5-alpha competent cells via heat shock and incubated on an orbital plate shaker for 1 h at 37°C shaking at 1000 rpm, then transferred to a deep-well plate containing 900 µl LB medium per well (total 1 ml) which was incubated at 37°C over night at the same shaker settings. On the next day, 5 µl cell culture were directly added to 45 µl PCR reaction mix (Supplementary Table S3, Thermo Fisher Phusion High-fidelity PCR-Kit). Therefore, the screened sequences lack sequence verification as they are derived from multiclonal cultures. Consequently, a small fraction of the sequences may be misassembled or contain unintended mutations. The used PCR protocol included an initial high-temperature step to first lyse cells, and then amplified the linear fragment for PURE expression (10 min 98°C, 30x(15 s 98°C, 15 s 60°C, 35 s 72°C), 5 min 72°C). Amplified DNA was purified using the QIAquick PCR Purification kit following the standard protocol and concentration was measured by NanoDrop. To use the DNA as input to the PUREdrop, DNA was diluted to 9 nM in 100µl in a 96 well plate. For the validation experiments, genes encoding for sfGFP and mCherry where amplified from a pCoofy vector using primers annealing to T7 promotor and terminator regions.

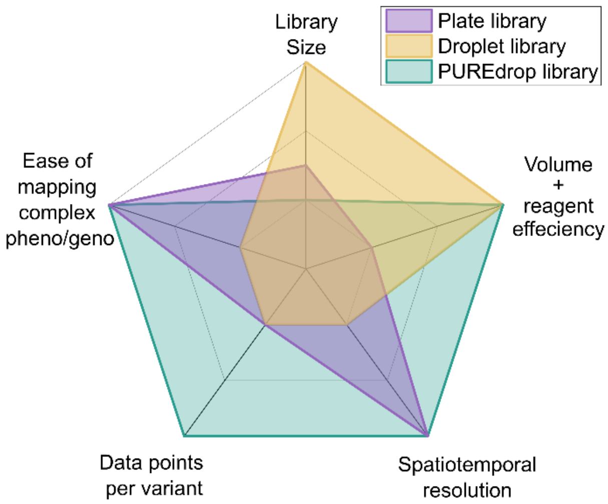


**Supplementary Fig. 1: Comparison between protein screening strategies grouped by library type.** The schematic highlights the differences in throughput, spatiotemporal resolution, ease of extracting genotype-phenotype linkage, and data richness across bulk well plate (arrayed), droplet-based sorting (pooled), and our PUREdrop (arrayed droplets) screening pipelines.

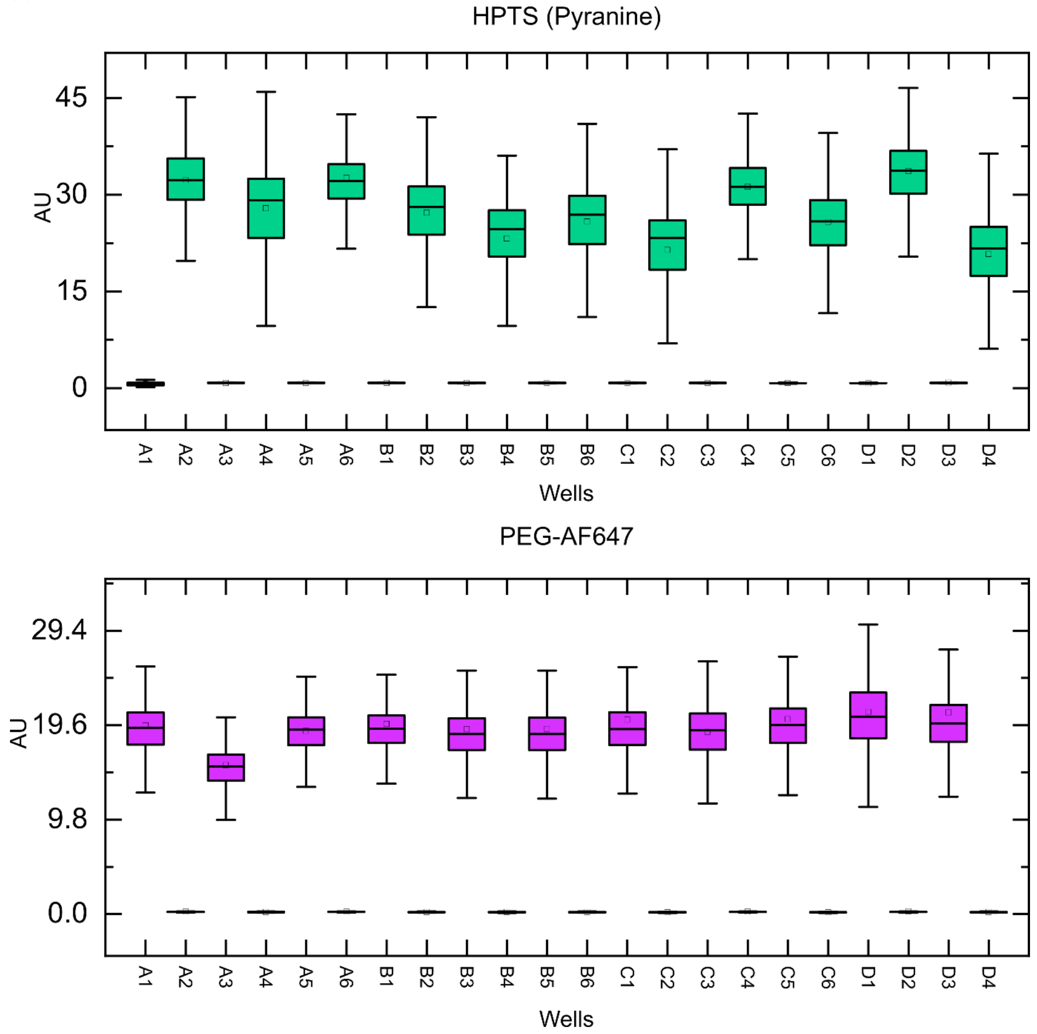


**Supplementary Fig. 2: Cross-contamination assay using alternating fluorescent tracers.** Schematic of the alternating input-well layout and corresponding box plots of fluorescence intensity distributions for droplet populations generated from wells containing Pyranine

(HPTS) or PEG-AF647. PEG-AF647 was prepared by conjugating 20 kDa PEG-azide with DBCO-AF647. Center lines indicate the median, boxes the interquartile range (IQR), and whiskers extend to 1.5 × IQR.

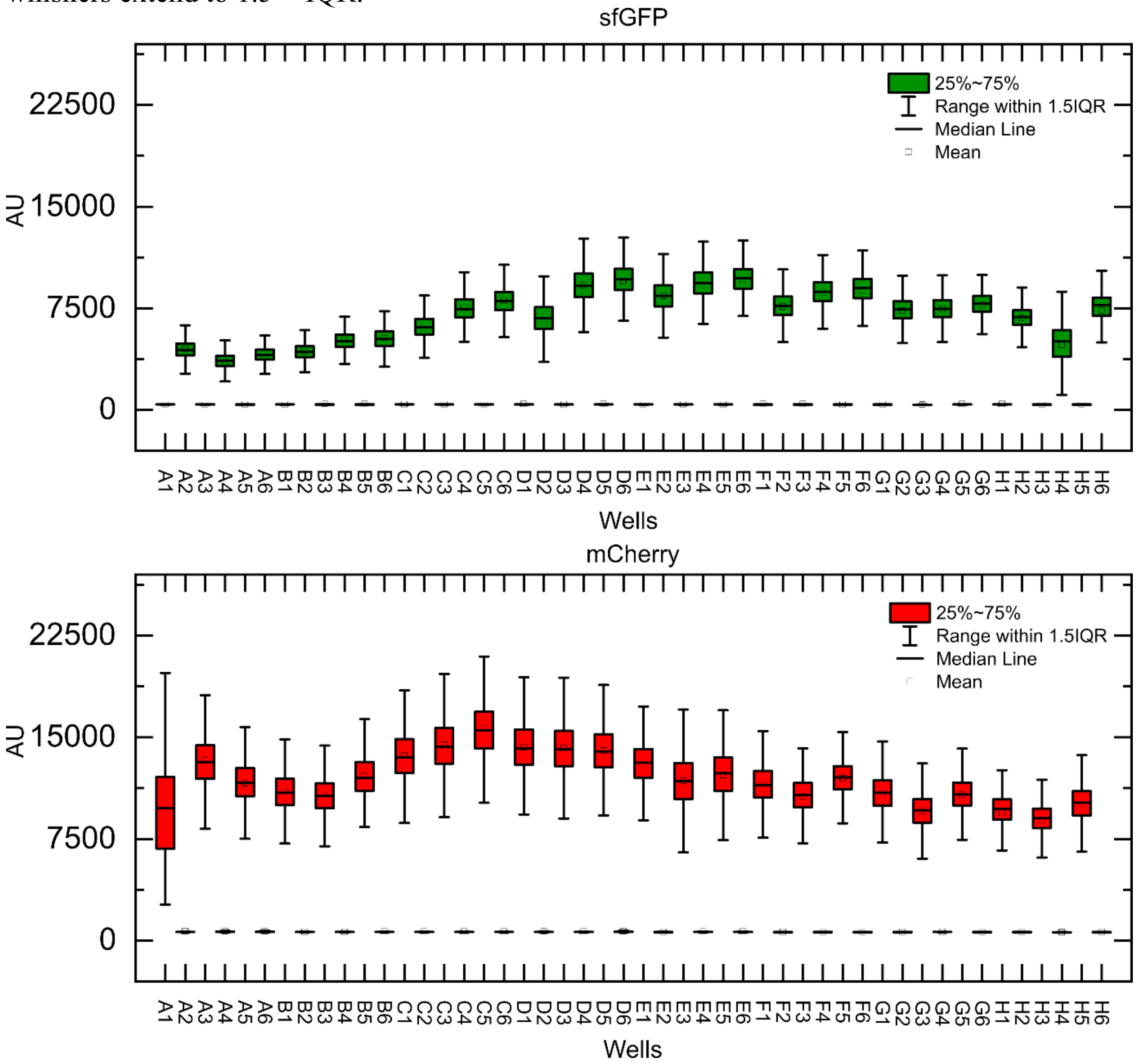


**Supplementary Fig. 3: Distribution of fluorescence intensities underlying Fig. 3b.** Non-normalized box plots showing the distribution of fluorescence intensities measured for all imaged droplets in each well after 15 h of expression for the indicated constructs. These data correspond to the well-averaged values summarized in Fig. 3b.

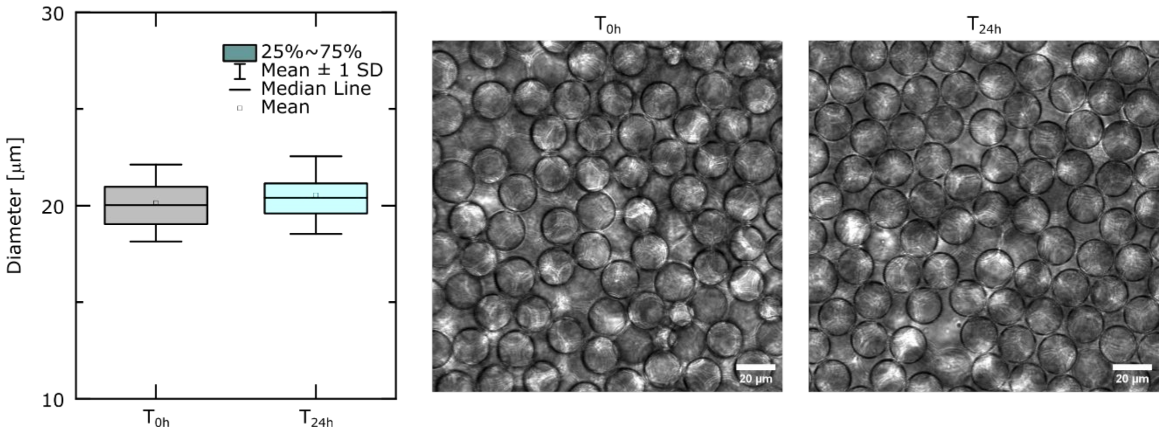


**Supplementary Fig. 4: Droplet stability during incubation at 30 °C.**
Representative images of droplets at the start of incubation ($n_0$ = 31,651) and after 24 h ($n_{24}$ = 65,535), demonstrating droplet stability over the full incubation period.

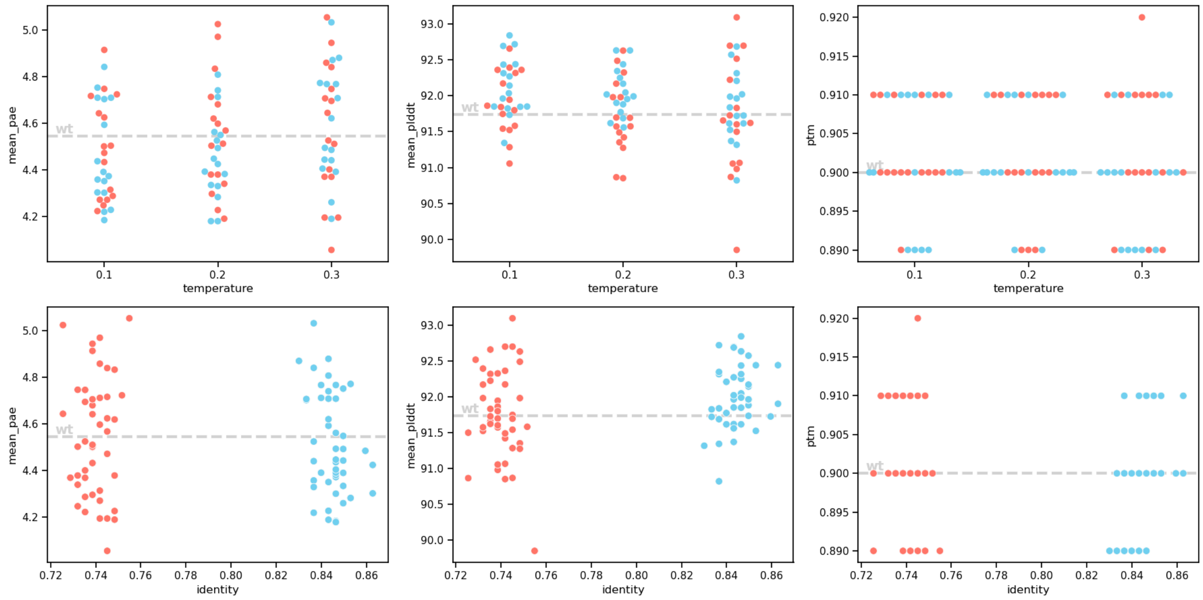


**Supplementary Fig. 5:** Confidence metrics of ColabFold prediction of the generated sequences. 70% conservation cutoff in blue, 50% conservation cutoff in red.

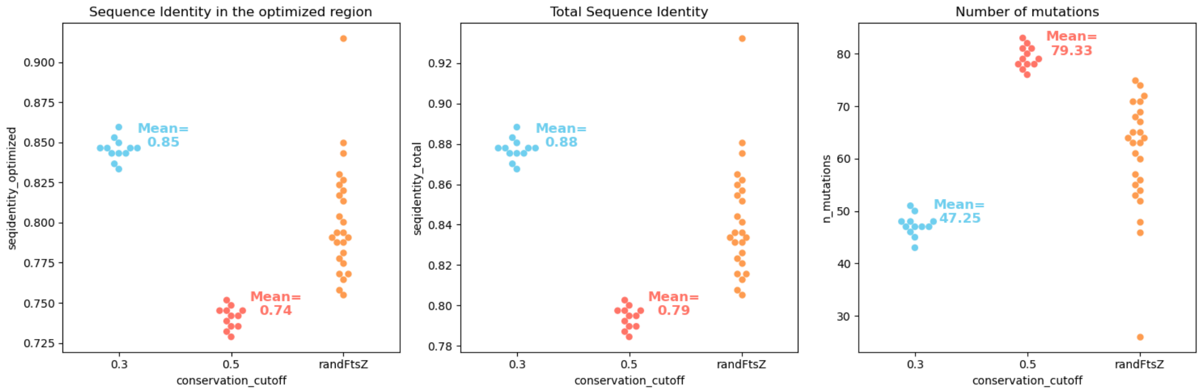


**Supplementary Fig. 6:** Sequence conservation of the 47 tested sequences. Left: Sequence identity to *E. coli* FtsZ when only taking the optimized region into account. Middle: Sequence identity to *E. coli* FtsZ when taking the full sequence into account, including unstructured parts that are always identical. Right: Total number of mutations.

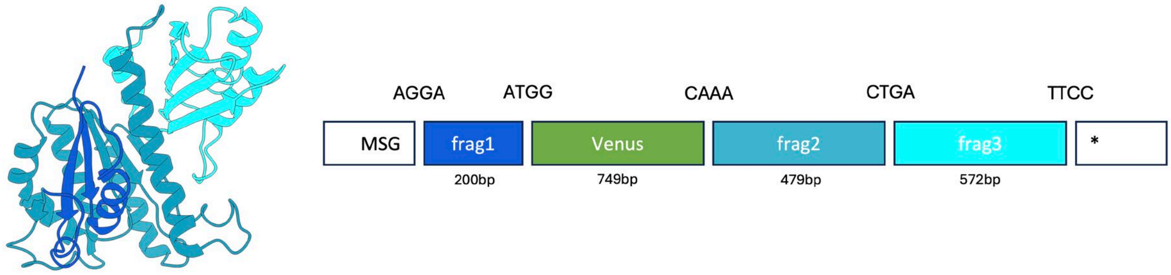


**Supplementary Fig 7:** Cloning scheme. All FtsZ variants and the wildtype were ordered in three fragments with the same overhangs. The fluorescent protein Venus is inserted between the first and second fragment, at position 55/56. The third fragment corresponds to the C-terminal domain.

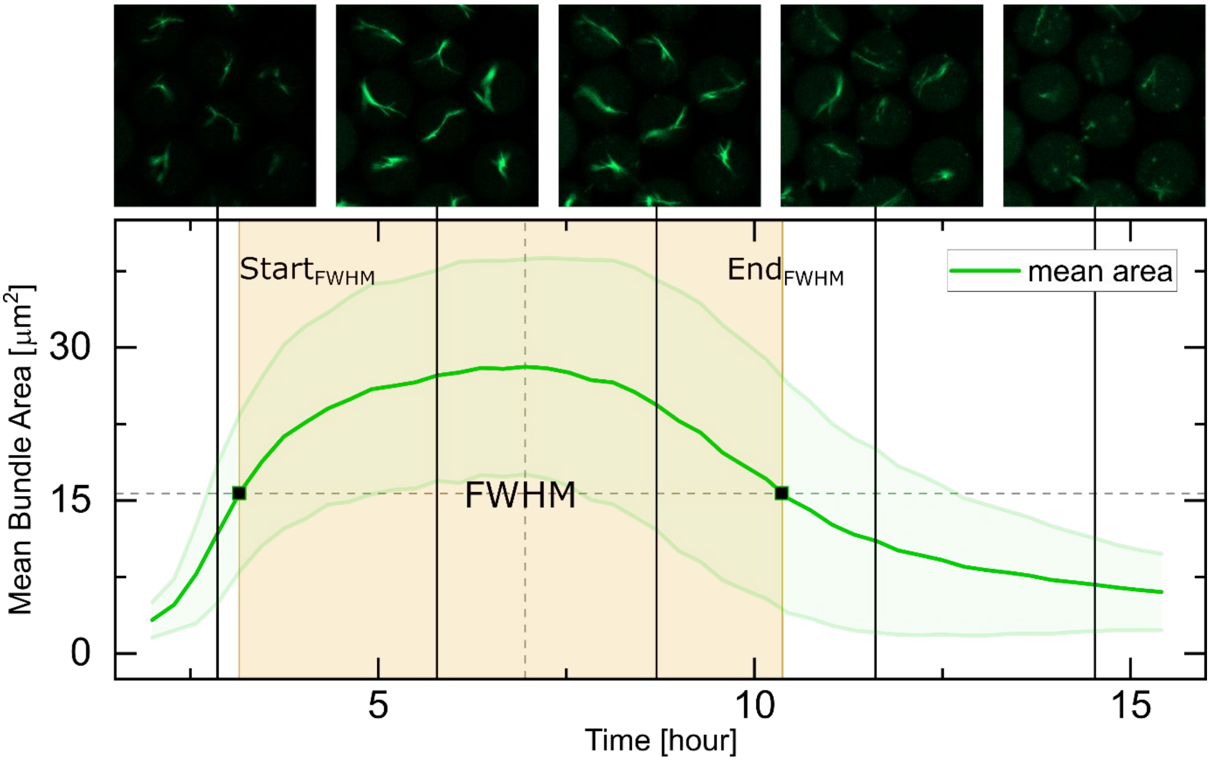


**Supplementary Fig. 8: Image collage of filament-bundle assembly and disassembly for the variant randFtzZv16 (G5).**
Representative montage showing bundle evolution at selected time points (2.9, 5.8, 8.7, 11.6 and 14.5 h), used to visually correlate changes in bundle area (FWHM) with the extracted bundle lifetimes presented in Fig. 6f–h.

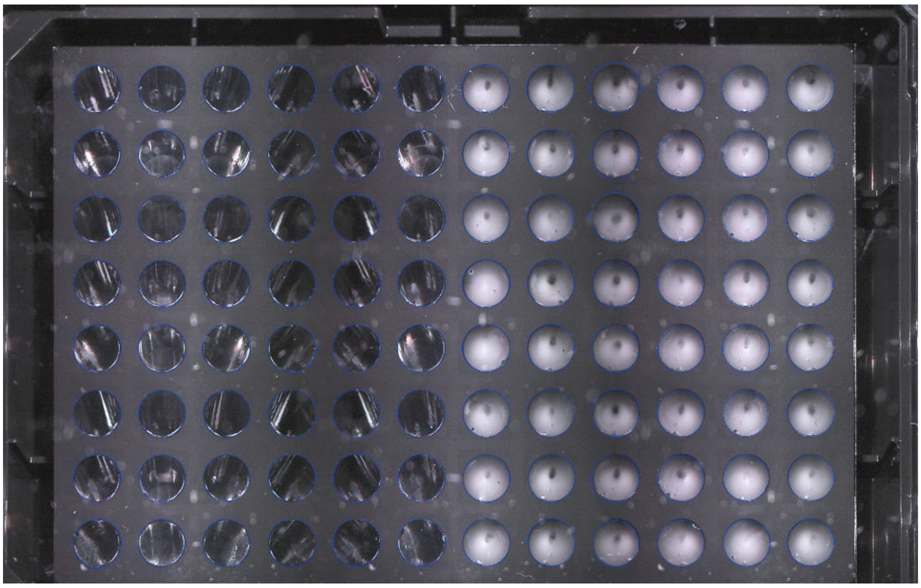

**Supplementary Fig. 9: Droplet packing in the output wells.**
Image of the 96-well plate collected after the PUREdrop sample-preparation run, showing the uniform packing of droplets within each well. The packed droplets give each well a turbidly white appearance.

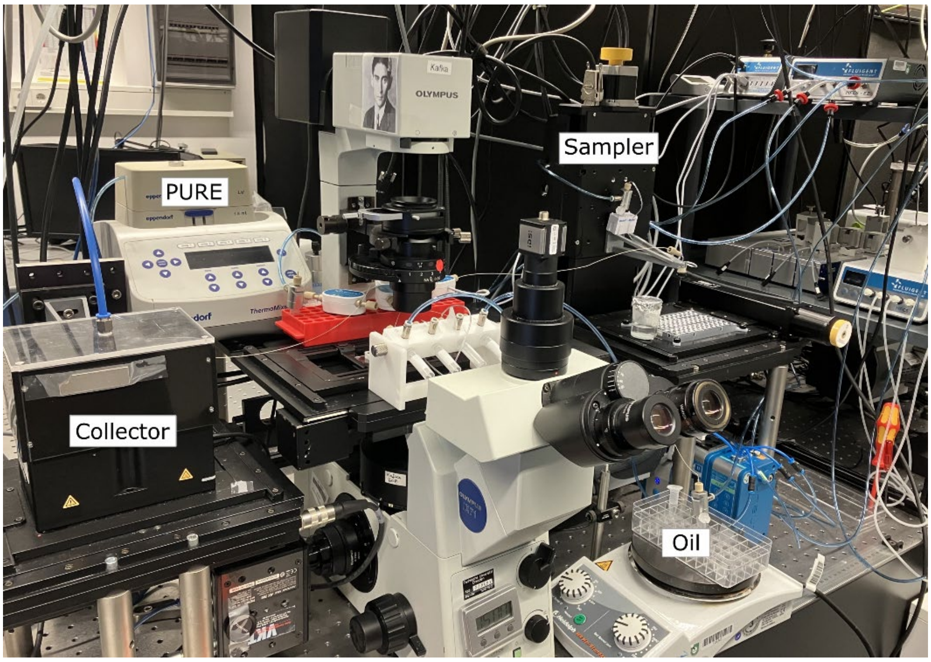


**Supplementary Fig. 10: Overview of the PUREdrop setup.**
Image of the assembled PUREdrop system, showing the physical placement of the autosampler and fraction collector relative to the microfluidic chip, which is mounted on the microscope.

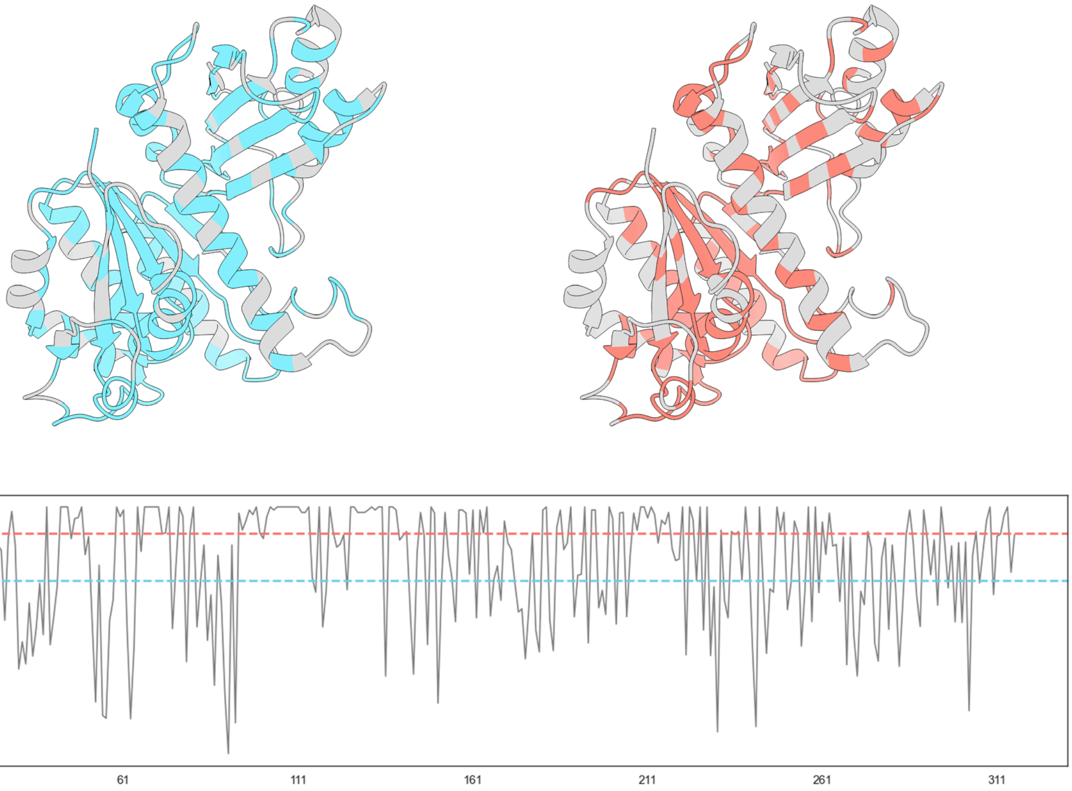


**Supplementary Fig. 11: Selection of conserved residues that were not re-designed.** A conservation score was calculated based on entropy (lower), and the most conserved 50% (red) or 70% (blue) of the residues were fixed. Conserved residues are mostly at the active site (bottom of protein structure) and in the core.

```
Kpneumoniae        1 AVIKVIGVGGGGGNAVEHMVRERIEGVEFFAVNTDAQALRKTAVGQTIQI      50
                     ||||||||||||||||||||||||||||||||||||||||||||||||||
Ecoli              1 AVIKVIGVGGGGGNAVEHMVRERIEGVEFFAVNTDAQALRKTAVGQTIQI      50

Kpneumoniae       51 GSGITKGLGAGANPEVGRNAADEDREALRAALDGADMVFIAAGMGGGTGT     100
                     ||||||||||||||||||||||||||:||||||:||||||||||||||||
Ecoli             51 GSGITKGLGAGANPEVGRNAADEDRDALRAALEGADMVFIAAGMGGGTGT     100

Kpneumoniae      101 GAAPVVAEVAKDLGILTVAVVTKPFNFEGKKRMAFAEQGITELSKHVDSL     150
                     ||||||||||||||||||||||||||||||||||||||||||||||||||
Ecoli            101 GAAPVVAEVAKDLGILTVAVVTKPFNFEGKKRMAFAEQGITELSKHVDSL     150

Kpneumoniae      151 ITIPNDKLLKVLGRGISLLDAFGAANDVLKGAVQGIAELITRPGLMNVDF     200
                     ||||||||||||||||||||||||||||||||||||||||||||||||||
Ecoli            151 ITIPNDKLLKVLGRGISLLDAFGAANDVLKGAVQGIAELITRPGLMNVDF     200

Kpneumoniae      201 ADVRTVMSEMGYAMMGSGVASGEDRAEEAAEMAISSPLLEDIDLSGARGV     250
                     ||||||||||||||||||||||||||||||||||||||||||||||||||
Ecoli            201 ADVRTVMSEMGYAMMGSGVASGEDRAEEAAEMAISSPLLEDIDLSGARGV     250

Kpneumoniae      251 LVNITAGFDLRLDEFETVGNTIRAFASDNATVVIGTSLDPDMNDELRVTV     300
                     ||||||||||||||||||||||||||||||||||||||||||||||||||
Ecoli            251 LVNITAGFDLRLDEFETVGNTIRAFASDNATVVIGTSLDPDMNDELRVTV     300

Kpneumoniae      301 VATGIG    306
                     ||||||
Ecoli            301 VATGIG    306
```

**Supplementary Fig. 12**: Sequence Alignment of *K. pneumoniae* and *E. coli* FtsZ. The two residues that differ are not conserved and are re-designed by ProteinMPNN.

A)

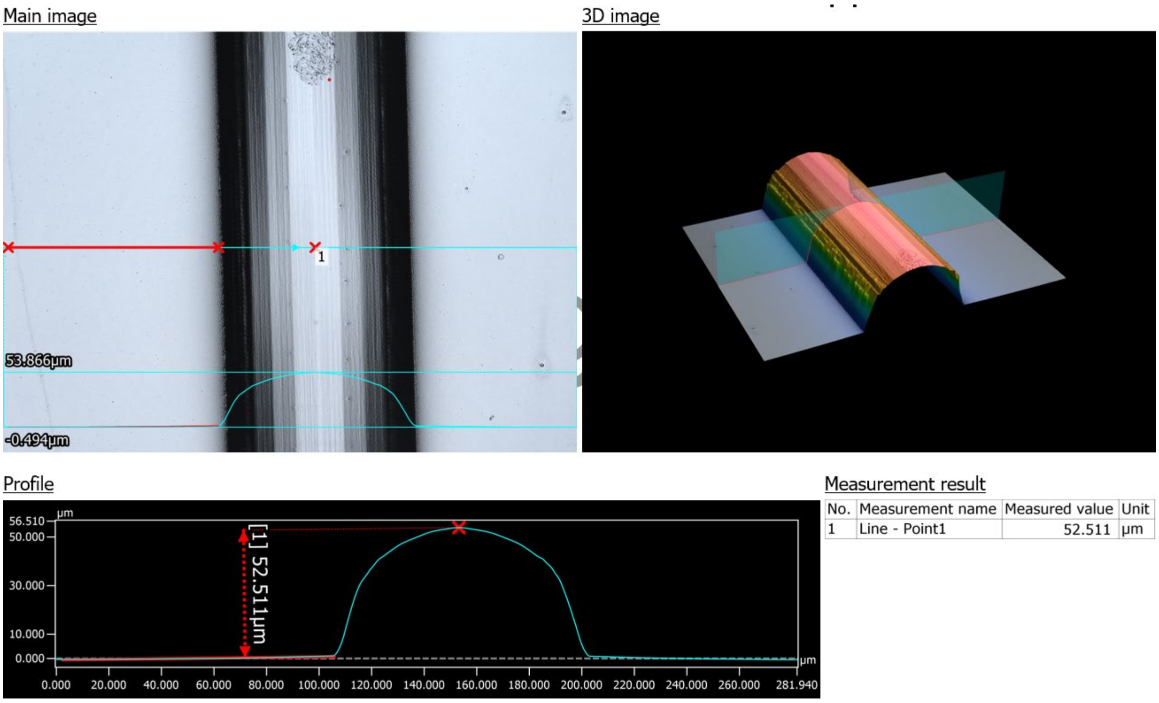


B)

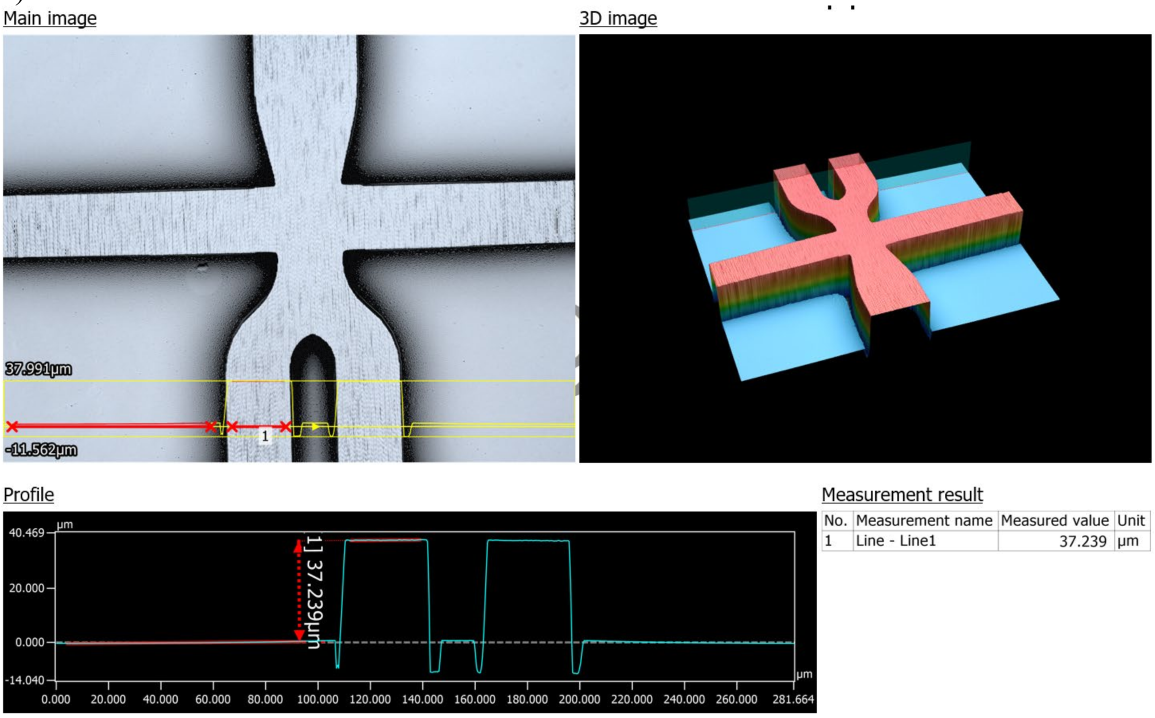


**Supplementary Fig. 13: Laser-profilometry measurements of the main-flow mold.** a, Height map and cross-sectional profile of the rounded channel geometry used for valve operation. b Height and profile of the five-way junction region of the chip, illustrating the transitions and feature uniformity across the intersection.

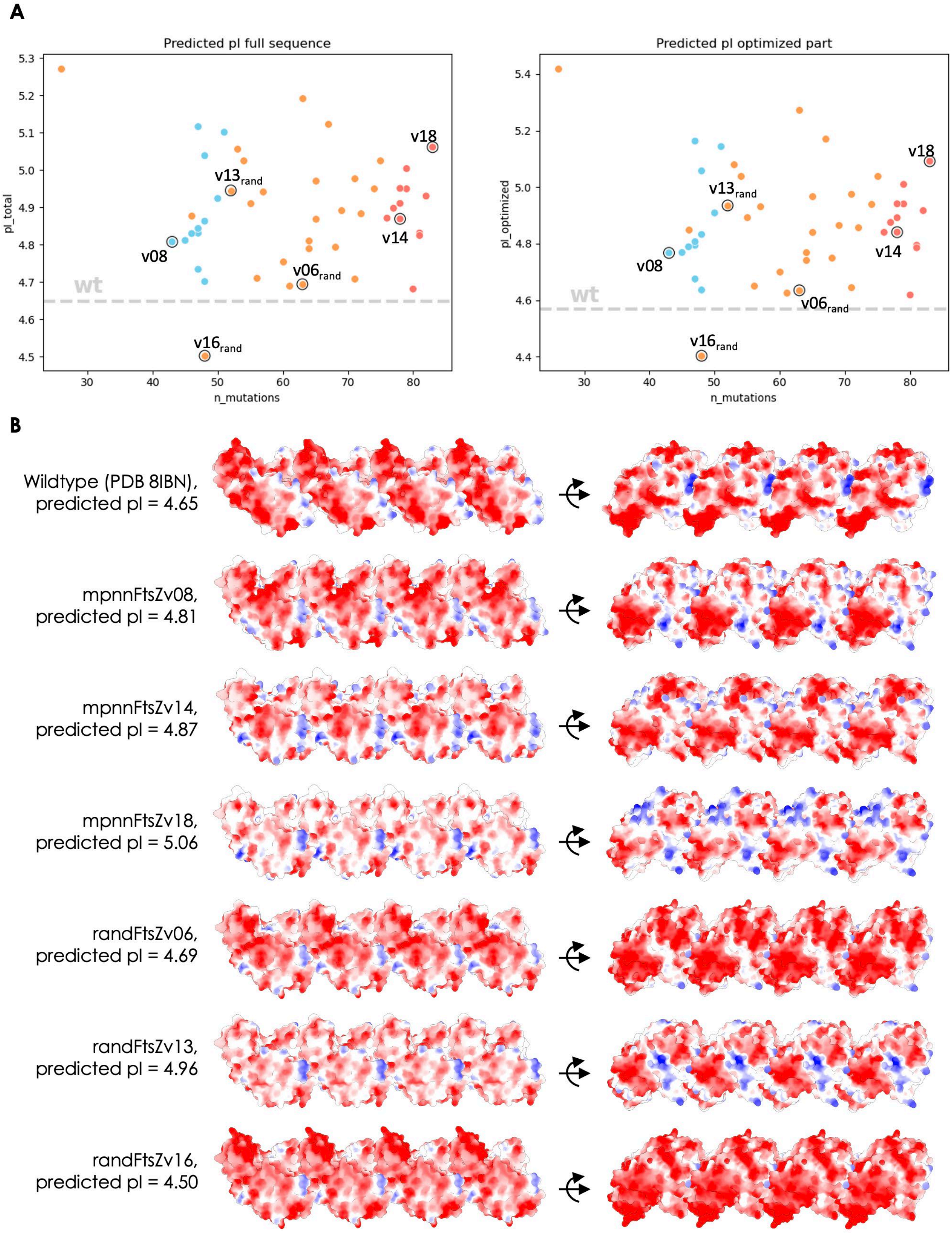


**Supplementary Fig. 14:** Effect of ProteinMPNN re-design on surface charge. **A:** Predicted pI of re-designed FtsZ variants for full protein (left) and only globular, re-designed part (right). For all but one variants, ProteinMPNN redesign decreased the overall charge. RandFtsZs in orange, 70% conservation cutoff in blue, 50% conservation cutoff in red. Variants functional in experiments highlighted with black circle. **B:** Surface colored by charge, calculated by ChimeraX. Red is negative, blue is positive. Isoelectric points were predicted with ProtParam in Biopython. While ProteinMPNN re-design reduces overall charge, it might introduce new local charged patches.

**Supplementary Table S1 | Sequence characteristics of FtsZ variants.** Sequence ID indicates the relationship of each construct to the wild-type (WT) FtsZ. The number of mutations reflects total amino acid substitutions relative to the WT sequence. The isoelectric point (pI) represents the theoretical pH at which the protein carries no net charge (surface charge for each variant hit is depicted in Supplementary Fig. 14). For shuffled re-designed sequences (rand), the composition column specifies the origin of each of the three constituent fragments, which are derived from different parent sequences. Fragments from non-functional variants are underlined.

Table 1: Synthetic FtsZ Hits

| Well | Name | Seq_ID | N_mutations | pI_total | Composition |
|---|---|---|---|---|---|
| D3 | ecFtsZ | 1.00 | 0 | 4.65 | ec/ec/ec |
| B2 | mpnnFtsZv08 | 0.89 | 43 | 4.81 | v08/v08/v08 |
| C2 | mpnnFtsZv14 | 0.80 | 78 | 4.87 | v14/v14/v14 |
| C6 | mpnnFtsZv18 | 0.78 | 83 | 5.06 | v18/v18/v18 |
| F1 | randFtsZv6 | 0.84 | 63 | 4.69 | v14/<u>v20</u>/v08 |
| G2 | randFtsZv13 | 0.86 | 52 | 4.95 | ec/v18/<u>v02</u> |
| G5 | randFtsZv16 | 0.88 | 48 | 4.50 | v18/<u>v16</u>/ec |

**Supplementary Table S2: Composition of a single GGA reaction**

| | |
|---|---|
| Fragment 1 (stock 20 ng/µl) | 0,075 µl |
| Fragment 2 (stock 40ng/µl) | 0,150 µl |
| Fragment 3 (stock 20 ng/µl) | 0,175 µl |
| Fragment 4 (stock 20 ng/µl) | 0,200 µl |
| Vector (stock 666 ng/µl) | 0,031 µl |
| H2O | 0,369 µl |
| 10x T4 Ligase Buffer (NEB M0202S) | 0,200 µl |
| BsaI-HFv2 (NEB R3733S) | 0,300 µl = 6 units / reaction |
| T4 Ligase (NEB M0202S) | 0,500 µl = 200 units / reaction |
| **TOTAL** | **2 µl** |

**Supplementary Table S3: The reaction mixture for cell lysis and FtsZ-gene variants amplification**

| | |
|---|---|
| Buffer GC (Thermo Fisher Phusion High-fidelity PCR-Kit) | 10 µl |
| dNTPs (stock 12,5mM) | 1 µl |
| Primer F (5'-CCCGCGAAATTAATACGACTCAC-3'; stock 50µM) | 0,25 µl |
| Primer R (5'-CAAAAAACCCCTCAAGACCCGT-3'; stock 50µM) | 0,25 µl |
| MgCl (stock 50mM) | 2 µl |
| DMSO (Thermo Fisher Phusion High-fidelity PCR-Kit) | 1 µl |
| Phusion Polymerase (stock 2 E/µl) | 0,5 µl |
| H2O | 30 µl |
| cells | 5 µl |
| **TOTAL** | **50 µl** |

**Supplementary Data 1:** Sequences of the wildtype and optimized FtsZ variants as tested (MSG- as first 3 residues due to cloning overhangs, wildtype has only M-)

>ecFtsZ

MSGFEPMELTNDAVIKVIGVGGGGGNAVEHMVRERIEGVEFFAVNTDAQALRKTAVGQTIQIGSGITKG
LGAGANPEVGRNAADEDRDALRAALEGADMVFIAAGMGGGTGTGAAPVVAEVAKDLGILTVAVVTKP
FNFEGKKRMAFAEQGITELSKHVDSLITIPNDKLLKVLGRGISLLDAFGAANDVLKGAVQGIAELITRPG
LMNVDFADVRTVMSEMGYAMMGSGVASGEDRAEEAAEMAISSPLLEDIDLSGARGVLVNITAGFDLRL
DEFETVGNTIRAFASDNATVVIGTSLDPDMNDELRVTVVATGIGMDKRPEITLVTNKQVQQPVMDRYQQ
HGMAPLTQEQKPVAKVVNDNAPQTAKEPDYLDIPAFLRKQAD
>mpnnFtsZv01

MSGFEPMELTNDAKIKVIGVGGGGGNAVDHMVGKELKGVDFVVVNTDAQALRKTAVGQTLQIGEELT
KGLGAGANPEVGRKAAEEDREKLREVLEGADMVFIAAGMGGGTGTGAAPVVAEVAKDLGILTVAVVT
KPFSFEGKKRLEFAEEGIEELSKVVDSLIEIPNDKLLKVLGKGISLLDAFGLANDVLRGAVEGIADLITRP
GLMNVDFADVRTVMSEMGRAMMGVGVAKGPNRAEEAAKAAITSPLLENIDLKGAKGVLVNITAGFDL
RLDEFEAVGNTIREFASDNATVVIGTSLDPGMGDELRVTVVATGIGMDKRPEITLVTNKQVQQPVMDRY
QQHGMAPLTQEQKPVAKVVNDNAPQTAKEPDYLDIPAFLRKQAD

>mpnnFtsZv02
MSGFEPMELTNDAKIKVIGVGGGGGNAVDHMVGKELKGVDFVVVNTDAQALRKTAVGQTLQIGEELT
KGLGAGANPEVGRKAAEEDRELLRKELEGADMVFIAAGMGGGTGTGAAPVVAKVAKDLGILTVAVVT
KPFEFEGKKRLEFAEKGIEELSKVVDSLIIIPNNKLLKVLGEGISLLDAFGLANDVLAGAVEGIADLITRPG
LMNVDFADVRTVMSEMGRAMMGVGVAKGPNRAEEAAKKAIESPLLENIDLEGAKGVLVNITAGFDLR
LDEFEAVGNTIRKFASDNATVVIGTSLEPGMGDELRVTVVATGIGMDKRPEITLVTNKQVQQPVMDRYQ
QHGMAPLTQEQKPVAKVVNDNAPQTAKEPDYLDIPAFLRKQAD

>mpnnFtsZv03
MSGFEPMELTNDAKIKVIGVGGGGGNAVDHMVGRELKGVDFIVVNTDAQALRKTAVGQTLQIGEELTK
GLGAGANPEVGRRAAEEDREKLEKALEGADMVFIAAGMGGGTGTGAAPVVAEVAKDLGILTVAVVTK
PFEFEGKKRLEFAEKGIEELSKVVDSLIIIPNNKLLKVLGDGISLLDAFGLANDVLRGAVEGIADLITRPGL
MNVDFADVRTVMSEMGRAMMGVGVAKGPNRAEEAAKAAITSPLLEDIDLKGAKGVLVNITAGFDLRL
DEFEAVGNTIREFASDNATVVIGTSLVPGMGDELRVTVVATGIGMDKRPEITLVTNKQVQQPVMDRYQQ
HGMAPLTQEQKPVAKVVNDNAPQTAKEPDYLDIPAFLRKQAD

>mpnnFtsZv04
MSGFEPMELTNDAKIKVIGVGGGGGNAVDHMVGRELDGVEFIVVNTDAQALRKTAVGQTLQIGEELTK
GLGAGANPEVGRRAALEDRAKLEEALTGADMVFIAAGMGGGTGTGAAPVVAEVAKDLGILTVAVVTK
PFEFEGKKRLEFAEKGIEELSKVVDSLIVIPNEKLLKVLGDGISLLDAFGLANDVLWGAVTGIADLITRPG
LMNVDFADVRTVMSEMGYAMMGVGVARGENRAEEAAQKAITSPLLENIDLEGAKGVLVNITSGFDLR
LDEFEAVGNTIRKFASDNATVVIGTSLEPGMGDELRVTVVATGIGMDKRPEITLVTNKQVQQPVMDRYQ
QHGMAPLTQEQKPVAKVVNDNAPQTAKEPDYLDIPAFLRKQAD

>mpnnFtsZv05
MSGFEPMELTNDAKIKVIGVGGGGGNAVDHMVGRELKGVDFVVVNTDAQALRKTAVGQTLQIGENLT
KGLGAGANPEVGRRAAEEDRELLAKALAGADMVFIAAGMGGGTGTGAAPVVAEVAKDLGILTVAVVT
KPFEFEGKKRLEFAEKGIEELSKVVDSLIVIPNNKLLKVLGNGISLLDAFGLANDVLAGAVDGIAELITRP
GLMNVDFADVRTVMSEMGRAMMGVGVAKGENRAEEAAKKAIESPLLENIDLKGAKGVLVNITAGFDL
RLDEFEAVGNTIRKFASDNATVVIGTSLVPGMGDELRVTVVATGIGMDKRPEITLVTNKQVQQPVMDRY
QQHGMAPLTQEQKPVAKVVNDNAPQTAKEPDYLDIPAFLRKQAD

>mpnnFtsZv06
MSGFEPMELTNDAKIKVIGVGGGGGNAVDHMVGRELDGVEFIVVNTDAQALRKTAVGQTLQIGEELTK
GLGAGANPEVGRKAALEDRAKLEEALTGADMVFIAAGMGGGTGTGAAPVVAEVAKDLGILTVAVVTK
PFSFEGKKRLEFAEKGIEELSKVVDSLIEIPNDKLLKVLGDGISLLDAFGLANDVLWGAVTGIADLITRPG
LMNVDFADVRTVMSEMGKAMMGVGVARGENRAEEAAQKAITSPLLENIDLEGAKGVLVNITAGFDLR

LDEFEAVGNTIRKFASDNATVVIGTSLEPGMGDELRVTVVATGIGMDKRPEITLVTNKQVQQPVMDRYQ
QHGMAPLTQEQKPVAKVVNDNAPQTAKEPDYLDIPAFLRKQAD

>mpnnFtsZv07
MSGFEPMELTNDAKIKVIGVGGGGGNAVDHMVGKELTGVDFVVVNTDAQALRKTAVGQTLQIGTELT
KGLGAGANPEVGRRAAEEDIELLRKALEGADMVFIAAGMGGGTGTGAAPVVAKVAKDLGILTVAVVT
KPFEFEGKKRLEFAEKGIEELSKVVDSLIVIPNNKLLKVLGEGISLLDAFGLANDVLAGAVQGIADLITRP
GLMNVDFADVRTVMSEMGRAMMGVGVAVGPNRAEEAAEKAISSPLLENIDLEGAKGVLVNITAGFDL
RLDEFEAVGNTIRKFASDNATVVIGTSLDPGMGDELRVTVVATGIGMDKRPEITLVTNKQVQQPVMDRY
QQHGMAPLTQEQKPVAKVVNDNAPQTAKEPDYLDIPAFLRKQAD

>mpnnFtsZv08
MSGFEPMELTNDAKIKVIGVGGGGGNAVDHMVGEELEGVEFIVVNTDAQALRKTAVGQTLQIGEELTK
GLGAGANPEVGRKAALEDREKLREALAGADMVFIAAGMGGGTGTGAAPVVAEVAKDLGILTVAVVTK
PFEFEGKKRLEFAEKGIEELSKVVDSLIIIPNNKLLKVLGKGISLLDAFGLANDVLRGAVRGIADLITRPGL
MNVDFADVRTVMSEMGRAMMGVGVARGPNRAEEAARKAIESPLLEDIDLEGARGVLVNITAGFDLRL
DEFEAVGNTIREFASDNATVVIGTSLDPEMGDELRVTVVATGIGMDKRPEITLVTNKQVQQPVMDRYQQ
HGMAPLTQEQKPVAKVVNDNAPQTAKEPDYLDIPAFLRKQAD

>mpnnFtsZv09
MSGFEPMELTNDAKIKVIGVGGGGGNAVDHMVGKELEGVDFVVVNTDAQALRKTAVGQTLQIGEELT
KGLGAGANPEVGRKAAEEDIELLRKALEGADMVFIAAGMGGGTGTGAAPVVAKVAKDLGILTVAVVT
KPFEFEGKKRLEFAEKGIEELSKVVDSLIVIPNNKLLKVLGEGISLLDAFGLANDVLAGAVQGIADLITRP
GLMNVDFADVRTVMSEMGRAMMGVGVAKGPNRAEEAAEKAISSPLLENIDLEGAKGVLVNITAGFDL
RLDEFEAVGNTIRKFASDNATVVIGTSLDPGMGDELRVTVVATGIGMDKRPEITLVTNKQVQQPVMDRY
QQHGMAPLTQEQKPVAKVVNDNAPQTAKEPDYLDIPAFLRKQAD

>mpnnFtsZv10
MSGFEPMELTNDAKIKVIGVGGGGGNAVDHMVGRELKGVDFIVVNTDAQALRKTAVGQTLQIGEELTK
GLGAGANPEVGRRAAEEDREKLRAALEGADMVFIAAGMGGGTGTGAAPVVAEVAKDLGILTVAVVTK
PFEFEGKKRLEFAEKGIEELSKVVDSLIIIPNNKLLKVLGKGISLLDAFGLANDVLRGAVEGIADLITRPGL
MNVDFADVRTVMSEMGRAMMGVGVAKGPNRAEEAAKKAITSPLLENIDLKGAKGVLVNITAGFDLRL
DEFEAVGNTIREFASDNATVVIGTSLVPGMGDELRVTVVATGIGMDKRPEITLVTNKQVQQPVMDRYQQ
HGMAPLTQEQKPVAKVVNDNAPQTAKEPDYLDIPAFLRKQAD

>mpnnFtsZv11
MSGFEPMELTNDAKIKVIGVGGGGGNAVDHMVGKDLKDVDFVVVNTDAQALRKTAVGQTLQIGEALT
KGLGAGANPEVGRKAAEEDREKLREVLKGADMVFIAAGMGGGTGTGAAPVVAEVAKDLGILTVAVVT
KPFSFEGKKRREFAEKGIEELSKVVDSLIIIPNDKLLKVLGNGISLLDAFGLANDVLRGAVEGIQDIITRPG
LMNVDFADVRTVMSEMGRAMMGVGVAKGPNRAEEAAKKAIESPLLENIDLNGAKGVLVNITAGFDLR
LDEFEAVGNTIRKFASDNATVVIGTSLDPGMGDELRVTVVATGIGMDKRPEITLVTNKQVQQPVMDRYQ
QHGMAPLTQEQKPVAKVVNDNAPQTAKEPDYLDIPAFLRKQAD

>mpnnFtsZv12
MSGFEPMELTNDARIKVIGVGGGGGNAVDHMVGKETKGVEFIVVNTDAQALRKTAVGQTLQIGENLTK
GLGAGANPEVGRRAAEEDRALLEAALAGADMVFIAAGMGGGTGTGAAPVVAEVAKDLGILTVAVVTK
PFEFEGKKRLEFAEKGIEELSKVVDSLIEIPNSKLLKVLGNGISLLDAFGLANDVLAGAVEGIADAITRPG
LMNVDFADVRTVMSEMGKAMMGVGVAKGPNRAEEAAKKAISSPLLEDIDLKGAKGVLVNITSGFDLR
LDEFEAVGNTIREFASDNATVVIGTSLDPGMGDELRVTVVATGIGMDKRPEITLVTNKQVQQPVMDRYQ
QHGMAPLTQEQKPVAKVVNDNAPQTAKEPDYLDIPAFLRKQAD

>mpnnFtsZv13
MSGFEPMELTNDKKIVVVGVGGGGGNALDHMAGKELKGITFIVVNTDAQALRKTAVGQTLQIGRELTG
GLGAGANPEIGRRAAEADREKLRAALRGADMVFIAAGMGGGTGTGAAPVVAEVAKELGILTVAVVTK
PFLFEGKKRQEFAEEGIEELREVVDSLIIIPNNTLLKVNGEGISLLDAFGLANDRLYEAVTGISDLITRPGL
MNVDFADVKTVMSAKGRAMIGVGRAEGEGRAEAAAREALTDPLLENPDLKGAKGVLVNITAGFDLRL
DEFEAIGNLIRSFASDNATVVIGTDIVPDEPGELRVTVVATGFGMDKRPEITLVTNKQVQQPVMDRYQQH
GMAPLTQEQKPVAKVVNDNAPQTAKEPDYLDIPAFLRKQAD

>mpnnFtsZv14

MSGFEPMELTNDKKIVVVGVGGGGGNALDHMAGKELEGVDFIVVNTDAQALRKTAVGQTLQIGRDLT
NGLGAGANPEVGREAAEKDRELLRKALEGADMVFIAAGMGGGTGTGAAPVVAEVAKELGILTVAVVT
KPFEFEGKKRLEFAEKGIKELEDVVDSLIEIPNSTLLKYKGDGISLLDAFGLANDRLSDAVYGILRLITRP
GLMNVDFADVKTVMASKGRAMIGVGRASGPNRAEAAAEKALTQPLLEDLDLAGAKGVLVNIESGFDL
RLDEFEAVGNRIREFASDNATVVIGTTIDPEKGGELTVTVVATGFGMDKRPEITLVTNKQVQQPVMDRY
QQHGMAPLTQEQKPVAKVVNDNAPQTAKEPDYLDIPAFLRKQAD

>mpnnFtsZv15
MSGFEPMELTNDKKILVVGVGGGGGNALDHMAGQPLEGIEFLVVNTDAQALRKTAVGQTLQIGSDLTQ
GLGAGANPEIGRKAAEQDLAKIRAALEGADMVFIAAGMGGGTGTGAAPVVAKAAKEKGILTVAVVTK
PFEFEGKKRLEFAEKGIEELKDVVDSLIVIPNSTLLKVNGEGISLLDAFGLANDRLRDAVTGIANLITRPG
LMNVDFADVKTVMSAKGRAMIGTGVATGPDRAEAAAKAALQDPLLENPDLEGAKGVLVNITAGFDLR
LDEFEAIGNLIRKFASDNATVVIGTVIEPDVPGELRVTVVATGFGMDKRPEITLVTNKQVQQPVMDRYQQ
HGMAPLTQEQKPVAKVVNDNAPQTAKEPDYLDIPAFLRKQAD

>mpnnFtsZv16
MSGFEPMELTNDKKILVVGVGGGGGNALDHMAGKENKGITFVVVNTDAQALRKTAVGQTLQIGEALT
GGLGAGANPEVGREAALVDRELLEKALEGADMVFIAAGMGGGTGTGAAPVVAEVAKEKGILTVAVVT
KPFEFEGKKRLEFAEKGIEELREVVDSLIEIPNSTLLKVRGDGISLLDAFGLANKRLEDAVYGIADLITRP
GLMNVDFADVKTVMAAKGRAMIGTGRAEGPGRAEAAAEAALTDPLLENPDLEGAKGVLVNITAGFDL
RLDEFEAVGNRIREFASDNATVVIGTTIDPGRGPELTVTVVATGFGMDKRPEITLVTNKQVQQPVMDRYQ
QHGMAPLTQEQKPVAKVVNDNAPQTAKEPDYLDIPAFLRKQAD

>mpnnFtsZv17
MSGFEPMELTNDKKIVVVGVGGGGGNALDHMAGKELKDITFIVVNTDAQALRKTAVGQTLQIGRELTG
GLGAGANPEIGRRAAEADREKLRAALEGADMVFIAAGMGGGTGTGAAPVVAEVAKELGILTVAVVTKP
FEFEGKKRLEFAEKGIEELREVVDSLIIIPNNTLLKVNGEGISLLDAFGLANDRLYDAVTGIADLITRPGL
MNVDFADVKTVMSAKGRAMIGTGVASGPNRAEAAAREALTDPLLENPDLEGAKGVLVNITAGFDLRL
DEFEAIGNLIRKFASDNATVVIGTDIDPERPGELRVTVVATGFGMDKRPEITLVTNKQVQQPVMDRYQQH
GMAPLTQEQKPVAKVVNDNAPQTAKEPDYLDIPAFLRKQAD

>mpnnFtsZv18
MSGFEPMELTNDREILVVGVGGGGGNALDHMAGQPVEGVRFLVVNTDAQALRKTAVGQTLQIGEELT
GGLGAGANPELGRAAAEADRAKLEAALQGADMVFIAAGMGGGTGTGAAPVVAEVAKKLGILTVAVV
TKPFVFEGKKRLEFAEEGIELLKNVVDSIIIIPNNTLLKANGKGISLLDAFGLANEALRNAVTGIAELITRP
GLMNVDFADVKTVMKSKGFAMIGTGRATGPNRAEAAARAALTQPLLENPDLNGAKGVLVNIASGFDL
RLDEFEAVGNLIREFASDNATVVIGTSITPGEGPELRVTVVATGLGMDKRPEITLVTNKQVQQPVMDRYQ
QHGMAPLTQEQKPVAKVVNDNAPQTAKEPDYLDIPAFLRKQAD

>mpnnFtsZv19
MSGFEPMELTNDKKILVVGVGGGGGNALDHMAGQPLEGIEFLVVNTDAQALRKTAVGQTLQIGEDLTG
GLGAGANPEIGRLAAERDRAKLAAALEGADMVFIAAGMGGGTGTGAAPVVAEVAKEKGILTVAVVTK
PFEFEGKKRLEFAEKGIEELKDVVDSLIEIPNSTLLKVNGKGISLLDAFGLANDRLRDAVTGIANLITRPG
LMNVDFADVKTVMSAKGRAMIGTGVATGENRAEAAAKAALTDPLLENPDLEGAKGVLVNITAGFDLR
LDEFEAIGNLIRKFASDNATVVIGTTIDPDKPGELTVTVVATGFGMDKRPEITLVTNKQVQQPVMDRYQQ
HGMAPLTQEQKPVAKVVNDNAPQTAKEPDYLDIPAFLRKQAD

>mpnnFtsZv20
MSGFEPMELTNDKKILVVGVGGGGGNALDHMAGKELKGITFIAVNTDAQALRKTAVGQTLQIGEELTG
GLGAGANPEIGREAALRDREKLEKALEGADMVFIAAGMGGGTGTGAAPVVAEVAKEKGILTVAVVTKP
FEFEGKKRLEFAEEGIELLKEVVDSLIEIPNNTLLKVNGKGISLLDAFGLANKRLEDAVTGIADLITRPGL
MNVDFADVKTVMAAKGRAMIGTGVAEGENRAEKAAEAALTDPLLENPDLKGAKGVLVNITAGFDLRL
DEFEAIGNKIREFASDNATVVIGTDIIPGRGPELTVTVVATGFGMDKRPEITLVTNKQVQQPVMDRYQQH
GMAPLTQEQKPVAKVVNDNAPQTAKEPDYLDIPAFLRKQAD

>mpnnFtsZv21
MSGFEPMELTNDKEIVVVGVGGGGGNALDHMAGKELKGIRFIVVNTDAQALRKTAVGQTLQIGEDLTG
GLGAGANPEIGRLAAERDRAKLAAALEGADMVFIAAGMGGGTGTGAAPVVAEVAKEKGILTVAVVTK
PFEFEGKKRLEFAEKGIEELRDVVDSLIEIPNNTLLKVNGKGISLLDAFGLANDRLEDAVTGIADLITRPG
LMNVDFADVKTVMAAKGRAMIGTGVAKGENRAEEAAKKALTDPLLENPDLKGAKGVLVNITAGFDL

RLDEFEAVGNLIREFASDNATVVIGTNIIPGEDGELRVTVVATGFGMDKRPEITLVTNKQVQQPVMDRYQ
QHGMAPLTQEQKPVAKVVNDNAPQTAKEPDYLDIPAFLRKQAD

>mpnnFtsZv22
MSGFEPMELTNDKKIVVVGVGGGGGNALDHMAGKELKGIDFIVVNTDAQALRKTAVGQTLQIGEELT
NGLGAGANPEIGREAAEKDREKLREALEGADMVFIAAGMGGGTGTGAAPVVAEVAKEKGILTVAVVT
KPFEFEGKKRLEFAEKGIKELEDVVDSLIEIPNSTLLKVNGKGISLLDAFGLANDRLREAVEGIARLITRP
GLMNVDFADVKTVMAAKGRAMIGVGVAEGPNRAEEAAEKALTQPLLENPDLKGAKGVLVNIEAGFDL
RLDEFEAVGNKIREFASDNATVVIGTDIIPGRGPELRVTVVATGFGMDKRPEITLVTNKQVQQPVMDRYQ
QHGMAPLTQEQKPVAKVVNDNAPQTAKEPDYLDIPAFLRKQAD

>mpnnFtsZv23
MSGFEPMELTNDREILVVGVGGGGGNALDHMAGEENKGIRFVVVNTDAQALRKTAVGQTLQIGEALT
GGLGAGANPEVGREAALVDRELLEKELEGADMVFIAAGMGGGTGTGAAPVVAEVAKEKGILTVAVVT
KPFEFEGKKRLEFAEKGIEELRDVVDSLIEIPNSTLLKVRGDGISLLDAFGLANDRLRDAVYGIARLITRP
GLMNVDFADVKTVMAAKGRAMIGTGVAEGPGRAEAAAEAALADPLLENPDLEGAKGVLVNIEAGFD
LRLDEFEAVGNRIREFASDNATVVIGTDIIPGEGPELRVTVVATGFGMDKRPEITLVTNKQVQQPVMDRY
QQHGMAPLTQEQKPVAKVVNDNAPQTAKEPDYLDIPAFLRKQAD

>mpnnFtsZv24
MSGFEPMELTNDSKIVVVGVGGGGGNALDHMAGKELKGITFIVVNTDAQALRKTAVGQTLQIGEELTG
GLGAGANPEIGREAAEADREKLEAALEGADMVFIAAGMGGGTGTGAAPVVAEVAKEKGILTVAVVTKP
FEFEGKKRLEFAEKGIEELREVVDSLIEIPNNTLLKVRGDGISLLDAFGLANDRLYDAVTGIADLITRPGL
MNVDFADVKTVMAAKGRAMIGTGVAEGPGRAEAAARAALTDPLLENPDLKGAKGVLVNIEAGFDLRL
DEFEAVGNLIRKFASDNATVVIGTDIVPGRPGELRVTVVATGFGMDKRPEITLVTNKQVQQPVMDRYQQ
HGMAPLTQEQKPVAKVVNDNAPQTAKEPDYLDIPAFLRKQAD

>randFtsZv1
MSGFEPMELTNDAKIKVIGVGGGGGNAVDHMVGRELKGVDFIVVNTDAQALRKTAVGQTLQIGEELTN
GLGAGANPEIGREAAEKDREKLREALEGADMVFIAAGMGGGTGTGAAPVVAEVAKEKGILTVAVVTKP
FEFEGKKRLEFAEKGIKELEDVVDSLIEIPNSTLLKVNGKGISLLDAFGLANDRLREAVEGIARLITRPGL
MNVDFADVRTVMSEMGKAMMGVGVAKGPNRAEEAAKKAISSPLLEDIDLKGAKGVLVNITSGFDLRL
DEFEAVGNTIREFASDNATVVIGTSLDPGMGDELRVTVVATGIGMDKRPEITLVTNKQVQQPVMDRYQQ
HGMAPLTQEQKPVAKVVNDNAPQTAKEPDYLDIPAFLRKQAD

>randFtsZv2
MSGFEPMELTNDKEIVVVGVGGGGGNALDHMAGKELKGIRFIVVNTDAQALRKTAVGQTLQIGEELTK
GLGAGANPEVGRKAALEDRAKLEEALTGADMVFIAAGMGGGTGTGAAPVVAEVAKDLGILTVAVVTK
PFSFEGKKRLEFAEKGIEELSKVVDSLIEIPNDKLLKVLGDGISLLDAFGLANDVLWGAVTGIADLITRPG
LMNVDFADVKTVMAAKGRAMIGTGVAEGPGRAEAAARAALTDPLLENPDLKGAKGVLVNIEAGFDLR
LDEFEAVGNLIRKFASDNATVVIGTDIVPGRPGELRVTVVATGFGMDKRPEITLVTNKQVQQPVMDRYQ
QHGMAPLTQEQKPVAKVVNDNAPQTAKEPDYLDIPAFLRKQAD

>randFtsZv3
MSGFEPMELTNDARIKVIGVGGGGGNAVDHMVGKETKGVEFIVVNTDAQALRKTAVGQTLQIGEALTK
GLGAGANPEVGRKAAEEDREKLREVLKGADMVFIAAGMGGGTGTGAAPVVAEVAKDLGILTVAVVTK
PFSFEGKKRREFAEKGIEELSKVVDSLIIIPNDKLLKVLGNGISLLDAFGLANDVLRGAVEGIQDIITRPGL
MNVDFADVKTVMAAKGRAMIGTGVAKGENRAEEAAKKALTDPLLENPDLKGAKGVLVNITAGFDLR
LDEFEAVGNLIREFASDNATVVIGTNIIPGEDGELRVTVVATGFGMDKRPEITLVTNKQVQQPVMDRYQQ
HGMAPLTQEQKPVAKVVNDNAPQTAKEPDYLDIPAFLRKQAD

>randFtsZv4
MSGFEPMELTNDAKIKVIGVGGGGGNAVDHMVGRELDGVEFIVVNTDAQALRKTAVGQTLQIGEDLTG
GLGAGANPEIGRLAAERDRAKLAAALEGADMVFIAAGMGGGTGTGAAPVVAEVAKEKGILTVAVVTK
PFEFEGKKRLEFAEKGIEELKDVVDSLIEIPNSTLLKVNGKGISLLDAFGLANDRLRDAVTGIANLITRPG
LMNVDFADVKTVMKSKGFAMIGTGRATGPNRAEAAARAALTQPLLENPDLNGAKGVLVNIASGFDLR
LDEFEAVGNLIREFASDNATVVIGTSITPGEGPELRVTVVATGLGMDKRPEITLVTNKQVQQPVMDRYQQ
HGMAPLTQEQKPVAKVVNDNAPQTAKEPDYLDIPAFLRKQAD

>randFtsZv5

MSGFEPMELTNDKKIVVVGVGGGGGNALDHMAGKELKGIDFIVVNTDAQALRKTAVGQTLQIGEELT
KGLGAGANPEVGRRAALEDRAKLEEALTGADMVFIAAGMGGGTGTGAAPVVAEVAKDLGILTVAVVT
KPFEFEGKKRLEFAEKGIEELSKVVDSLIVIPNEKLLKVLGDGISLLDAFGLANDVLWGAVTGIADLITRP
GLMNVDFADVKTVMAAKGRAMIGTGVAEGPGRAEAAAEAALADPLLENPDLEGAKGVLVNIEAGFD
LRLDEFEAVGNRIREFASDNATVVIGTDIIPGEGPELRVTVVATGFGMDKRPEITLVTNKQVQQPVMDRY
QQHGMAPLTQEQKPVAKVVNDNAPQTAKEPDYLDIPAFLRKQAD

>randFtsZv6
MSGFEPMELTNDKKIVVVGVGGGGGNALDHMAGKELEGVDFIVVNTDAQALRKTAVGQTLQIGEELT
GGLGAGANPEIGREAALRDREKLEKALEGADMVFIAAGMGGGTGTGAAPVVAEVAKEKGILTVAVVT
KPFEFEGKKRLEFAEEGIELLKEVVDSLIEIPNNTLLKVNGKGISLLDAFGLANKRLEDAVTGIADLITRP
GLMNVDFADVRTVMSEMGRAMMGVGVARGPNRAEEAARKAIESPLLEDIDLEGARGVLVNITAGFDL
RLDEFEAVGNTIREFASDNATVVIGTSLDPEMGDELRVTVVATGIGMDKRPEITLVTNKQVQQPVMDRY
QQHGMAPLTQEQKPVAKVVNDNAPQTAKEPDYLDIPAFLRKQAD

>randFtsZv7
MSGFEPMELTNDAKIKVIGVGGGGGNAVDHMVGKDLKDVDFVVVNTDAQALRKTAVGQTLQIGENLT
KGLGAGANPEVGRRAAEEDRALLEAALAGADMVFIAAGMGGGTGTGAAPVVAEVAKDLGILTVAVVT
KPFEFEGKKRLEFAEKGIEELSKVVDSLIEIPNSKLLKVLGNGISLLDAFGLANDVLAGAVEGIADAITRP
GLMNVDFADVKTVMSAKGRAMIGTGVASGPNRAEAAAREALTDPLLENPDLEGAKGVLVNITAGFDL
RLDEFEAIGNLIRKFASDNATVVIGTDIDPERPGELRVTVVATGFGMDKRPEITLVTNKQVQQPVMDRYQ
QHGMAPLTQEQKPVAKVVNDNAPQTAKEPDYLDIPAFLRKQAD

>randFtsZv8
MSGFEPMELTNDAKIKVIGVGGGGGNAVDHMVGRELDGVEFIVVNTDAQALRKTAVGQTLQIGRELTG
GLGAGANPEIGRRAAEADREKLRAALEGADMVFIAAGMGGGTGTGAAPVVAEVAKELGILTVAVVTKP
FEFEGKKRLEFAEKGIEELREVVDSLIIIPNNTLLKVNGEGISLLDAFGLANDRLYDAVTGIADLITRPGL
MNVDFADVRTVMSEMGRAMMGVGVAKGENRAEEAAKKAIESPLLENIDLKGAKGVLVNITAGFDLRL
DEFEAVGNTIRKFASDNATVVIGTSLVPGMGDELRVTVVATGIGMDKRPEITLVTNKQVQQPVMDRYQQ
HGMAPLTQEQKPVAKVVNDNAPQTAKEPDYLDIPAFLRKQAD

>randFtsZv9
MSGFEPMELTNDAKIKVIGVGGGGGNAVDHMVGKELTGVDFVVVNTDAQALRKTAVGQTLQIGRDLT
NGLGAGANPEVGREAAEKDRELLRKALEGADMVFIAAGMGGGTGTGAAPVVAEVAKELGILTVAVVT
KPFEFEGKKRLEFAEKGIKELEDVVDSLIEIPNSTLLKYKGDGISLLDAFGLANDRLSDAVYGILRLITRP
GLMNVDFADVKTVMSAKGRAMIGTGVATGPDRAEAAAKAALQDPLLENPDLEGAKGVLVNITAGFDL
RLDEFEAIGNLIRKFASDNATVVIGTVIEPDVPGELRVTVVATGFGMDKRPEITLVTNKQVQQPVMDRYQ
QHGMAPLTQEQKPVAKVVNDNAPQTAKEPDYLDIPAFLRKQAD

>randFtsZv10
MSGFEPMELTNDAKIKVIGVGGGGGNAVDHMVGRELKGVDFIVVNTDAQALRKTAVGQTLQIGRELT
GGLGAGANPEIGRRAAEADREKLRAALRGADMVFIAAGMGGGTGTGAAPVVAEVAKELGILTVAVVT
KPFLFEGKKRQEFAEEGIEELREVVDSLIIIPNNTLLKVNGEGISLLDAFGLANDRLYEAVTGISDLITRPG
LMNVDFADVKTVMAAKGRAMIGTGVAEGENRAEKAAEAALTDPLLENPDLKGAKGVLVNITAGFDLR
LDEFEAIGNKIREFASDNATVVIGTDIIPGRGPELTVTVVATGFGMDKRPEITLVTNKQVQQPVMDRYQQ
HGMAPLTQEQKPVAKVVNDNAPQTAKEPDYLDIPAFLRKQAD
>randFtsZv11
MSGFEPMELTNDKKIVVVGVGGGGGNALDHMAGKELKGITFIVVNTDAQALRKTAVGQTLQIGTELTK
GLGAGANPEVGRRAAEEDIELLRKALEGADMVFIAAGMGGGTGTGAAPVVAKVAKDLGILTVAVVTK
PFEFEGKKRLEFAEKGIEELSKVVDSLIVIPNNKLLKVLGEGISLLDAFGLANDVLAGAVQGIADLITRPG
LMNVDFADVRTVMSEMGRAMMGVGVAKGPNRAEEAAKAAITSPLLENIDLKGAKGVLVNITAGFDLR
LDEFEAVGNTIREFASDNATVVIGTSLDPGMGDELRVTVVATGIGMDKRPEITLVTNKQVQQPVMDRYQ
QHGMAPLTQEQKPVAKVVNDNAPQTAKEPDYLDIPAFLRKQAD

>randFtsZv12
MSGFEPMELTNDKKILVVGVGGGGGNALDHMAGKENKGITFVVVNTDAQALRKTAVGQTLQIGEDLT
GGLGAGANPEIGRLAAERDRAKLAAALEGADMVFIAAGMGGGTGTGAAPVVAEVAKEKGILTVAVVT
KPFEFEGKKRLEFAEKGIEELRDVVDSLIEIPNNTLLKVNGKGISLLDAFGLANDRLEDAVTGIADLITRP
GLMNVDFADVRTVMSEMGYAMMGVGVARGENRAEEAAQKAITSPLLENIDLEGAKGVLVNITSGFDL

RLDEFEAVGNTIRKFASDNATVVIGTSLEPGMGDELRVTVVATGIGMDKRPEITLVTNKQVQQPVMDRY
QQHGMAPLTQEQKPVAKVVNDNAPQTAKEPDYLDIPAFLRKQAD

>randFtsZv13
MSGFEPMELTNDAVIKVIGVGGGGGNAVEHMVRERIEGVEFFAVNTDAQALRKTAVGQTLQIGEELTGG
LGAGANPELGRAAAEADRAKLEAALQGADMVFIAAGMGGGTGTGAAPVVAEVAKKLGILTVAVVTKP
FVFEGKKRLEFAEEGIELLKNVVDSIIIIPNNTLLKANGKGISLLDAFGLANEALRNAVTGIAELITRPGLM
NVDFADVRTVMSEMGRAMMGVGVAKGPNRAEEAAKKAIESPLLENIDLEGAKGVLVNITAGFDLRLD
EFEAVGNTIRKFASDNATVVIGTSLEPGMGDELRVTVVATGIGMDKRPEITLVTNKQVQQPVMDRYQQH
GMAPLTQEQKPVAKVVNDNAPQTAKEPDYLDIPAFLRKQAD

>randFtsZv14
MSGFEPMELTNDREILVVGVGGGGGNALDHMAGEENKGIRFVVVNTDAQALRKTAVGQTLQIGSDLT
QGLGAGANPEIGRKAAEQDLAKIRAALEGADMVFIAAGMGGGTGTGAAPVVAKAAKEKGILTVAVVT
KPFEFEGKKRLEFAEKGIEELKDVVDSLIVIPNSTLLKVNGEGISLLDAFGLANDRLRDAVTGIANLITRP
GLMNVDFADVRTVMSEMGRAMMGVGVAKGPNRAEEAAKAAITSPLLEDIDLKGAKGVLVNITAGFDL
RLDEFEAVGNTIREFASDNATVVIGTSLVPGMGDELRVTVVATGIGMDKRPEITLVTNKQVQQPVMDRY
QQHGMAPLTQEQKPVAKVVNDNAPQTAKEPDYLDIPAFLRKQAD

>randFtsZv15
MSGFEPMELTNDAKIKVIGVGGGGGNAVDHMVGKELEGVDFVVVNTDAQALRKTAVGQTLQIGEELT
KGLGAGANPEVGRKAAEEDREKLREVLEGADMVFIAAGMGGGTGTGAAPVVAEVAKDLGILTVAVVT
KPFSFEGKKRLEFAEEGIEELSKVVDSLIEIPNDKLLKVLGKGISLLDAFGLANDVLRGAVEGIADLITRP
GLMNVDFADVKTVMSAKGRAMIGTGVATGENRAEAAAKAALTDPLLENPDLEGAKGVLVNITAGFDL
RLDEFEAIGNLIRKFASDNATVVIGTTIDPDKPGELTVTVVATGFGMDKRPEITLVTNKQVQQPVMDRYQ
QHGMAPLTQEQKPVAKVVNDNAPQTAKEPDYLDIPAFLRKQAD

>randFtsZv16
MSGFEPMELTNDREILVVGVGGGGGNALDHMAGQPVEGVRFLVVNTDAQALRKTAVGQTLQIGEALT
GGLGAGANPEVGREAALVDRELLEKALEGADMVFIAAGMGGGTGTGAAPVVAEVAKEKGILTVAVVT
KPFEFEGKKRLEFAEKGIEELREVVDSLIEIPNSTLLKVRGDGISLLDAFGLANKRLEDAVYGIADLITRP
GLMNVDFADVRTVMSEMGYAMMGSGVASGEDRAEEAAEMAISSPLLEDIDLSGARGVLVNITAGFDLR
LDEFETVGNTIRAFASDNATVVIGTSLDPDMNDELRVTVVATGIGMDKRPEITLVTNKQVQQPVMDRYQ
QHGMAPLTQEQKPVAKVVNDNAPQTAKEPDYLDIPAFLRKQAD

>randFtsZv17
MSGFEPMELTNDAKIKVIGVGGGGGNAVDHMVGKELKGVDFVVVNTDAQALRKTAVGQTLQIGEELT
KGLGAGANPEVGRKAAEEDRELLRKELEGADMVFIAAGMGGGTGTGAAPVVAKVAKDLGILTVAVVT
KPFEFEGKKRLEFAEKGIEELSKVVDSLIIIPNNKLLKVLGEGISLLDAFGLANDVLAGAVEGIADLITRPG
LMNVDFADVKTVMAAKGRAMIGTGRAEGPGRAEAAAEAALTDPLLENPDLEGAKGVLVNITAGFDLR
LDEFEAVGNRIREFASDNATVVIGTTIDPGRGPELTVTVVATGFGMDKRPEITLVTNKQVQQPVMDRYQ
QHGMAPLTQEQKPVAKVVNDNAPQTAKEPDYLDIPAFLRKQAD

>randFtsZv18
MSGFEPMELTNDKKIVVVGVGGGGGNALDHMAGKELKDITFIVVNTDAQALRKTAVGQTLQIGEELTK
GLGAGANPEVGRRAAEEDREKLRAALEGADMVFIAAGMGGGTGTGAAPVVAEVAKDLGILTVAVVTK
PFEFEGKKRLEFAEKGIEELSKVVDSLIIIPNNKLLKVLGKGISLLDAFGLANDVLRGAVEGIADLITRPGL
MNVDFADVRTVMSEMGRAMMGVGVAKGPNRAEEAAKKAIESPLLENIDLNGAKGVLVNITAGFDLRL
DEFEAVGNTIRKFASDNATVVIGTSLDPGMGDELRVTVVATGIGMDKRPEITLVTNKQVQQPVMDRYQQ
HGMAPLTQEQKPVAKVVNDNAPQTAKEPDYLDIPAFLRKQAD

>randFtsZv19
MSGFEPMELTNDAKIKVIGVGGGGGNAVDHMVGRELKGVDFVVVNTDAQALRKTAVGQTIQIGSGITK
GLGAGANPEVGRNAADEDRDALRAALEGADMVFIAAGMGGGTGTGAAPVVAEVAKDLGILTVAVVT
KPFNFEGKKRMAFAEQGITELSKHVDSLITIPNDKLLKVLGRGISLLDAFGAANDVLKGAVQGIAELITR
PGLMNVDFADVRTVMSEMGRAMMGVGVAKGPNRAEEAAKKAITSPLLENIDLKGAKGVLVNITAGFD
LRLDEFEAVGNTIREFASDNATVVIGTSLVPGMGDELRVTVVATGIGMDKRPEITLVTNKQVQQPVMDR
YQQHGMAPLTQEQKPVAKVVNDNAPQTAKEPDYLDIPAFLRKQAD

>randFtsZv20

MSGFEPMELTNDKKILVVGVGGGGGNALDHMAGKELKGITFIAVNTDAQALRKTAVGQTLQIGEELTG
GLGAGANPEIGREAAEADREKLEAALEGADMVFIAAGMGGGTGTGAAPVVAEVAKEKGILTVAVVTKP
FEFEGKKRLEFAEKGIEELREVVDSLIEIPNNTLLKVRGDGISLLDAFGLANDRLYDAVTGIADLITRPGL
MNVDFADVRTVMSEMGRAMMGVGVAVGPNRAEEAAEKAISSPLLENIDLEGAKGVLVNITAGFDLRL
DEFEAVGNTIRKFASDNATVVIGTSLDPGMGDELRVTVVATGIGMDKRPEITLVTNKQVQQPVMDRYQQ
HGMAPLTQEQKPVAKVVNDNAPQTAKEPDYLDIPAFLRKQAD

>randFtsZv21
MSGFEPMELTNDKKILVVGVGGGGGNALDHMAGQPLEGIEFLVVNTDAQALRKTAVGQTLQIGEELTK
GLGAGANPEVGRRAAEEDREKLEKALEGADMVFIAAGMGGGTGTGAAPVVAEVAKDLGILTVAVVTK
PFEFEGKKRLEFAEKGIEELSKVVDSLIIIPNNKLLKVLGDGISLLDAFGLANDVLRGAVEGIADLITRPGL
MNVDFADVKTVMASKGRAMIGVGRASGPNRAEAAAEKALTQPLLEDLDLAGAKGVLVNIESGFDLRL
DEFEAVGNRIREFASDNATVVIGTTIDPEKGGELTVTVVATGFGMDKRPEITLVTNKQVQQPVMDRYQQ
HGMAPLTQEQKPVAKVVNDNAPQTAKEPDYLDIPAFLRKQAD

>randFtsZv22
MSGFEPMELTNDSKIVVVGVGGGGGNALDHMAGKELKGITFIVVNTDAQALRKTAVGQTLQIGEELTK
GLGAGANPEVGRKAAEEDIELLRKALEGADMVFIAAGMGGGTGTGAAPVVAKVAKDLGILTVAVVTK
PFEFEGKKRLEFAEKGIEELSKVVDSLIVIPNNKLLKVLGEGISLLDAFGLANDVLAGAVQGIADLITRPG
LMNVDFADVKTVMSAKGRAMIGVGRAEGEGRAEAAAREALTDPLLENPDLKGAKGVLVNITAGFDLR
LDEFEAIGNLIRSFASDNATVVIGTDIVPDEPGELRVTVVATGFGMDKRPEITLVTNKQVQQPVMDRYQQ
HGMAPLTQEQKPVAKVVNDNAPQTAKEPDYLDIPAFLRKQAD

>randFtsZv23
MSGFEPMELTNDKKILVVGVGGGGGNALDHMAGQPLEGIEFLVVNTDAQALRKTAVGQTLQIGEELTK
GLGAGANPEVGRKAALEDREKLREALAGADMVFIAAGMGGGTGTGAAPVVAEVAKDLGILTVAVVTK
PFEFEGKKRLEFAEKGIEELSKVVDSLIIIPNNKLLKVLGKGISLLDAFGLANDVLRGAVRGIADLITRPGL
MNVDFADVKTVMAAKGRAMIGVGVAEGPNRAEEAAEKALTQPLLENPDLKGAKGVLVNIEAGFDLRL
DEFEAVGNKIREFASDNATVVIGTDIIPGRGPELRVTVVATGFGMDKRPEITLVTNKQVQQPVMDRYQQH
GMAPLTQEQKPVAKVVNDNAPQTAKEPDYLDIPAFLRKQAD

>FtsZoptG55VenusQ56
MFEPMELTNDAVIKVIGVGGGGGNAVEHMVRERIEGVEFFAVNTDAQALRKTAVGMVSKGEELFTGVV
PILVELDGDVNGHKFSVSGEGEGDATYGKLTLKLICTTGKLPVPWPTLVTTLGYGLQCFARYPDHMKQ
HDFFKSAMPEGYVQERTIFFKDDGNYKTRAEVKFEGDTLVNRIELKGIDFKEDGNILGHKLEYNYNSHN
VYITADKQKNGIKANFKIRHNIEDGGVQLADHYQQNTPIGDGPVLLPDNHYLSYQSALSKDPNEKRDH
MVLLEFVTAAGITLGMDELYKQTIQIGSGITKGLGAGANPEVGRNAADEDRDALRAALEGADMVFIAA
GMGGGTGTGAAPVVAEVAKDLGILTVAVVTKPFNFEGKKRMAFAEQGITELSKHVDSLITIPNDKLLKV
LGRGISLLDAFGAANDVLKGAVQGIAELITRPGLMNVDFADVRTVMSEMGYAMMGSGVASGEDRAEE
AAEMAISSPLLEDIDLSGARGVLVNITAGFDLRLDEFETVGNTIRAFASDNATVVIGTSLDPDMNDELRV
TVVATGIGMDKRPEITLVTNKQVQQPVMDRYQQHGMAPLTQEQKPVAKVVNDNAPQTAKEPDYLDIPA
FLRKQAD

>cytoFtsN DDEE
MAQRDYVRRSQPAPSDDEESTSRKKQRNLPAV

>cytoFtsN RAAK
MAQRDYVRRSQPAPSRAAKSTSRKKQRNLPAV

>cytoFtsN wt
MAQRDYVRRSQPAPSRRKKSTSRKKQRNLPAV

> KIL
MIAHHFGTDEIPRQCVTPGDYVLHEGRTYIASANNIKKRKLYIRNLTTKTFITDRMIKVFLGRDGLPVKA
ESW

> SlmA
MAEKQTAKRNRREEILQSLALMLESSDGSQRITTAKLAASVGVSEAALYRHFPSKTRMFDSLIEFIEDSLI
TRINLILKDEKDTTARLRLIVLLLLGFGERNPGLTRILTGHALMFEQDRLQGRINQLFERIEAQLRQVLRE
KRMREGEGYTTDETLLASQILAFCEGMLSRFVRSEFKYRPTDDFDARWPLIAAQLQ

>FtsAopt
MIKATDRKLVVGLEIGTAKVAALVGEVLPDGMVNIIGVGSCPSRGMDKGGVNDLESVVKCVQRAIDQA
ELMADCQISSVYLALSGKHISCQNEIGMVPISEEEVTQEDVENVVHTAKSVRVRDEHRVLHVIPQEYAID
YQEGIKNPVGLSGVRMQAKVHLITCHNDMAKNIVKAVERCGLKVDQLIFAGLASSYSVLTEDERELGV
CVVDIGGGTMDIAVYTGGALRHTKVIPYAGNVVTSDIAYAFGTPPSDAEAIKVRHGCALGSIVGKDESV
EVPSVGGRPPRSLQRQTLAEVIEPRYTELLNLVNEEILQLQEKLRQQGVKHHLAAGIVLTGGAAQIEGLA
ACAQRVFHTQVRIGAPLNITGLTDYAQEPYYSTAVGLLHYGKESHLNGEAEVEKRVTASVGSWIKRLNS
WLRKEF

> FtsAopt R268W
MIKATDRKLVVGLEIGTAKVAALVGEVLPDGMVNIIGVGSCPSRGMDKGGVNDLESVVKCVQRAIDQA
ELMADCQISSVYLALSGKHISCQNEIGMVPISEEEVTQEDVENVVHTAKSVRVRDEHRVLHVIPQEYAID
YQEGIKNPVGLSGVRMQAKVHLITCHNDMAKNIVKAVERCGLKVDQLIFAGLASSYSVLTEDERELGV
CVVDIGGGTMDIAVYTGGALRHTKVIPYAGNVVTSDIAYAFGTPPSDAEAIKVRHGCALGSIVGKDESV
EVPSVGGRPPWSLQRQTLAEVIEPRYTELLNLVNEEILQLQEKLRQQGVKHHLAAGIVLTGGAAQIEGL
AACAQRVFHTQVRIGAPLNITGLTDYAQEPYYSTAVGLLHYGKESHLNGEAEVEKRVTASVGSWIKRLN
SWLRKEF

> ZipA
MMQDLRLILIIVGAIAIIALLVHGFWTSRKERSSMFRDRPLKRMKSKRDDDSYDEDVEDDEGVGEVRVH
RVNHAPANAQEHEAARPSPQHQYQPPYASAQPRQPVQQPPEAQVPPQHAPHPAQPVQQPAYQPQPEQP
LQQPVSPQVAPAPQPVHSAPQPAQQAFQPAEPVAAPQPEPVAEPAPVMDKPKRKEAVIIMNVAAHHGSEL
NGELLLNSIQQAGFIFGDMNIYHRHLSPDGSGPALFSLANMVKPGTFDPEMKDFTTPGVTIFMQVPSYG
DELQNFKLMLQSAQHIADEVGGVVLDDQRRMMTPQKLREYQDIIREVKDANA

> ClpX
MTDKRKDGSGKLLYCSFCGKSQHEVRKLIAGPSVYICDECVDLCNDIIREEIKEVAPHRERSALPTPHEIR
NHLDDYVIGQEQAKKVLAVAVYNHYKRLRNGDTSNGVELGKSNILLIGPTGSGKTLLAETLARLLDVPF
TMADATTLTEAGYVGEDVENIIQKLLQKCDYDVQKAQRGIVYIDEIDKISRKSDNPSITRDVSGEGVQQ
ALLKLIEGTVAAVPPQGGRKHPQQEFLQVDTSKILFICGGAFAGLDKVISHRVETGSGIGFGATVKAKSD
KASEGELLAQVEPEDLIKFGLIPEFIGRLPVVATLNELSEEALIQILKEPKNALTKQYQALFNLEGVDLEFR
DEALDAIAKKAMARKTGARGLRSIVEAALLDTMYDLPSMEDVEKVVIDESVIDGQSEPLLIYGKPEAQ
QASGE

> ClpP
MPYSGERDNFAPHMALVPMVIEQTSRGERSFDIYSRLLKERVIFLTGQVEDHMANLIVAQMLFLEAENPE
KDIYLYINSPGGVITAGMSIYDTMQFIKPDVSTICMGQAASMGAFLLTAGAKGKRFCLPNSRVMIHQPLG
GYQGQATDIEIHAREILKVKGRMNELMALHTGQSLEQIERDTERDRFLSAPEAVEYGLVDSILTHRN

>His-mCherry
MKHHHHHHSAGLEVLFQGPMVSKGEEDNMAIIKEFMRFKVHMEGSVNGHEFEIEGEGEGRPYEGTQT
AKLKVTKGGPLPFAWDILSPQFMYGSKAYVKHPADIPDYLKLSFPEGFKWERVMNFEDGGVVTVTQDS
SLQDGEFIYKVKLRGTNFPSDGPVMQKKTMGWEASSERMYPEDGALKGEIKQRLKLKDGGHYDAEV
KTTYKAKKPVQLPGAYNVNIKLDITSHNEDYTIVEQYERAEGRHSTGGMDELYK

>mCherry2-MinC
MVSKGEEDNMAIIKEFMRFKVHMEGSVNGHEFEIEGEGEGRPYEGTQTAKLKVTKGGPLPFAWDILSP
QFMYGSKAYVKHPADIPDYLKLSFPEGFNWERVMNFEDGGVVTVTQDSSLQDGEFIYKVKLRGTNFPS
DGPVMQCRTMGWEASTERMYPEDGALKGEIKQRLKLKDGGHYDAEVKTTYKAKKPVQLPGAYNVDI
KLDILSHNEDYTIVEQYERAEGRHSTGGMDELYKGAAGEFSNTPIELKGSSFTLSVVHLHEAEPKVIHQ
ALEDKIAQAPAFLKHAPVVLNVSALEDPVNWSAMHKAVSATGLRVIGVSGCKDAQLKAEIEKMGLPIL
TEGKEKAPRPAPTPQAPAQNTTPVTKTRLIDTPVRSGQRIYAPQCDLIVTSHVSAGAELIADGNIHVYGM
MRGRALAGASGDRETQIFCTNLMAELVSIAGEYWLSDQIPAEFYGKAARLQLVENALTVQPLN

>sfGFP
MVSKGEEDNMAIIKEFMRFKVHMEGSVNGHEFEIEGEGEGRPYEGTQTAKLKVTKGGPLPFAWDILSP
QFMYGSKAYVKHPADIPDYLKLSFPEGFNWERVMNFEDGGVVTVTQDSSLQDGEFIYKVKLRGTNFPS
DGPVMQCRTMGWEASTERMYPEDGALKGEIKQRLKLKDGGHYDAEVKTTYKAKKPVQLPGAYNVDI
KLDILSHNEDYTIVEQYERAEGRHSTGGMDELYKGAAGEFSNTPIELKGSSFTLSVVHLHEAEPKVIHQ

ALEDKIAQAPAFLKHAPVVLNVSALEDPVNWSAMHKAVSATGLRVIGVSGCKDAQLKAEIEKMGLPIL
TEGKEKAPRPAPTPQAPAQNTTPVTKTRLIDTPVRSGQRIYAPQCDLIVTSHVSAGAELIADGNIHVYGM
MRGRALAGASGDRETQIFCTNLMAELVSIAGEYWLSDQIPAEFYGKAARLQLVENALTVQPLN


Desai, S. P., Freeman, D. M., & Voldman, J. (2009). Plastic masters - Rigid templates for soft lithography. *Lab on a Chip*, *9*(11), 1631–1637. https://doi.org/10.1039/b822081f

Gernhardt, M., Frisch, H., Welle, A., Jones, R., Wegener, M., Blasco, E., & Barner-Kowollik, C. (2020). Multi-material 3D microstructures with photochemically adaptive mechanical properties. *Journal of Materials Chemistry C*, *8*(32), 10993–11000. https://doi.org/10.1039/d0tc02751k

Qian, J., Milles, L. F., Wicky, B. I. M., Ragotte, R. J., Motmaen, A., Borst, A. J., Skotheim, R., Ols, S., Coventry, B., Li, X., Kibler, R. D., Goreshnik, I., Expòsit, M., Loré, K., Stewart, L., & Baker, D. (2025). Accelerating protein design by scaling experimental characterization. In *bioRxiv*. Cold Spring Harbor Laboratory. https://doi.org/10.1101/2025.08.05.668824
